# Annexin A2 modulates phospholipid membrane composition upstream of Arp2 to control angiogenic sprout initiation

**DOI:** 10.1101/2022.07.07.498997

**Authors:** Timothy M. Sveeggen, Colette A. Abbey, Rebecca L. Smith, Kayla J. Bayless

**Affiliations:** Department of Molecular and Cellular Medicine, Texas A&M Health Science Center, TX, USA; Interdisciplinary Graduate Program in Genetics, Texas A&M University, TX, USA

**Keywords:** Key Terms Actin-Related Protein 2, Adherens Junctions, Annexin A2, Cholesterol, Cytoskeleton, Endothelial Cells, Lipidomics, Membrane Lipids, Phosphatidylcholines, Phospholipids

## Abstract

The intersection of protein and lipid biology is of growing importance for understanding how cells address structural challenges during adhesion and migration. While protein complexes engaged with the cytoskeleton play a vital role, support from the phospholipid membrane is crucial for directing localization and assembly of key protein complexes. During angiogenesis, it is well observed that dramatic cellular remodeling is necessary for endothelial cells to shift from a stable monolayer to invasive structures. However, the molecular dynamics between lipids and proteins during endothelial invasion are not defined. Here, we utilized cell culture, immunofluorescence, and lipidomic analyses to identify a novel role for the membrane binding protein Annexin A2 (ANXA2) in modulating the composition of specific membrane lipids necessary for cortical F-actin organization and adherens junction stabilization. In the absence of ANXA2, there is disorganized cortical F-actin, reduced junctional Arp2, excess sprout initiation, and ultimately failed sprout maturation. Further, we observed reduced filipin III labeling of membrane cholesterol in cells with reduced ANXA2, suggesting there is an alteration in phospholipid membrane dynamics. Lipidomic analyses reveal that 42 lipid species are altered with loss of ANXA2, including an accumulation of phosphatidylcholine (16:0_16:0). We find that supplementation of phosphatidylcholine (16:0_16:0) in wild-type endothelial cells mimics the ANXA2 knock-down phenotype, indicating that ANXA2 regulates the phospholipid membrane upstream of Arp2 recruitment and organization of cortical F-actin. Altogether these data indicate a novel role for ANXA2, and show that proper lipid modulation is a critical component of endothelial sprouting.

**Summary Statement:** Annexin A2 modulates composition of select phospholipid species in endothelial cells needed for F-actin organization and Arp2 recruitment to endothelial adherens junctions. These events simultaneously temper sprout initiation and support sprout maturation to maintain the integrity of sprouting structures during angiogenesis

## Introduction

The formation of new blood vessels from pre-existing structures creates an exceptional problem for endothelial cells, which are responsible for maintaining the vascular barrier. During angiogenesis, cell-cell junctions maintain the endothelial barrier and remodel to allow endothelial cells to move collectively during the outgrowth of new sprouts.^1, 2^ To accomplish this balance, specialized intracellular machinery must coordinate events at the junction that allow endothelial cells to both initiate sprouting and maintain cell-cell contacts.^3–7^ However, the mechanisms responsible for this regulation are not fully understood.

Angiogenesis is driven by a variety of signals, which can be generalized as enhancing or opposing barrier formation. An example is vascular endothelial growth factor (VEGF), which loosens the barrier between endothelial cells, leading to increased vessel permeability and vascular development.^8, 9^ Another potent growth factor, basic fibroblast growth factor (bFGF), will stabilize the endothelial barrier at low levels, yet increases vessel permeability in a dose-dependent manner at higher concentrations.^10–14^ Lastly, sphingosine 1-phosphate (S1P) is an abundant plasma lysosphingolipid secreted by activated platelets,^15^ which greatly enhances barrier stability. As plasma levels of S1P are maintained through incorporation into red blood cells and high density lipoproteins,^16, 17^ endothelial cells will consistently preserve a stable vascular barrier.^18–20^ Despite opposing effects on barrier formation between growth factors (VEGF and bFGF) and S1P, these agonists synergize to enhance endothelial sprouting in multiple models.^11, 21, 22^ One explanation for this observation is that the dynamic process of angiogenesis is optimized by a careful balance of signals that loosen junctions to initiate sprouting, then stabilize junctions as new multicellular structures form into a vessel with sufficient barrier integrity.^4^

Transitioning from a stable monolayer to an invasive sprout requires coordination between transmembrane protein complexes and the actin cytoskeleton. During this transition, the actin cytoskeleton must actively remodel to minimize stress on the cell.^23, 24^ Adaptor proteins such as Annexin A2 (ANXA2) are ideal candidates to minimize stress at the cytoskeletal-membrane interface, through binding to both F-actin and phospholipids. ANXA2 heavily localizes wherever dynamic cytoskeletal-membrane structures form,^25–29^ which correlates with its increased expression in cell types that are motile, complex in shape, and/or stress bearing, such as macrophages, fibroblasts, epithelial cells, and endothelial cells.^30, 31^ While ANXA2 is known to support endothelial junctional maintenance, barrier integrity, and sprout formation, the precise mechanisms by which ANXA2 supports these processes are unclear.^32–34^

It has long been argued that a critical function of ANXA2 is regulation of the phospholipid membrane, which directs protein localization. Traditionally, membrane lipid biology has broadly focused on lipid class ratios or levels of saturation, however, it is now appreciated that particular lipid species influence cellular functions,^35, 36^ and minor lipid species can drive protein localization to shape cell behaviors.^37^ Lipid-dependent signaling events involving lipoproteins, lysophospholipids, and phosphatidylinositol phosphates have been studied during angiogenesis,^38, 39^ but how endothelial cells alter membrane lipid composition when transitioning from a stable monolayer to an invasive structure is not understood, as the dynamic nature of angiogenesis likely requires equally dynamic shifts in the membrane itself.

In this study, we identify a new mechanism of function for ANXA2 in angiogenesis. In the absence of ANXA2, cortical actin and adherens junctions are disorganized and there is failed recruitment of Arp2, resulting in immediate excess sprout initiation, but ultimately failed sprout maturation. These events are explained in part by reduced labelling of cholesterol by filipin III, which suggests cholesterol presentation within the membrane is affected by loss of ANXA2.

Lipidomic analyses show ANXA2 is required for proper levels of 42 lipid species. Notably, loss of ANXA2 results in accumulation of phosphatidylcholine (PC) (16:0_16:0), also known as dipalmitoylphosphatidylcholine (DPPC). Treatment of human umbilical vein endothelial cells (HUVEC) with DPPC mimics the shANXA2 phenotype, leading to disruption of Arp2 recruitment, enhanced sprout initiation, and inhibition of sprout maturation. Altogether, the data shown here reveal a novel role for ANXA2 in S1P-mediated junctional stabilization and lipid homeostasis; ANXA2 maintains composition of the plasma membrane, which limits DPPC accumulation and excessive sprout initiation, while allowing for Arp2 recruitment. This ultimately enables maturation into a multicellular sprouting structure, which is crucial for angiogenesis.

## Materials and Methods

### Cell Culture

Authenticated human umbilical vein endothelial cells (HUVEC) obtained from Lonza (cat. C2517A, CC-2517) were cultured on 1 mg/ml gelatin-coated flasks prior to use for experiments at passage 3 – 6. Growth medium previously described^40^ consisted of M199 supplemented with 0.4 mg/ml bovine hypothalamic extract (Pel-Freeze Biologicals), 100 µg/ml heparin, 10% fetal bovine serum (FBS) (Lonza), penicillin and streptomycin. Human embryonic kidney 293 cells (HEK 293FT) obtained from Lonza were cultured on flasks coated with 20 μg/ml collagen type I. Growth medium previously described ^34^ consisted of Dulbecco’s modified eagle medium supplemented with 10% FBS, and antibiotics. HEK 293 used were within passages 3 – 14.

### Making lentivirus

For each shRNA construct, 1250 ng plasmid DNA was transfected into 1×10^6^ HEK 293FT cells with 1250 ng pLP1, pLP2, and pLP/VSVG packaging plasmids in a 25 cm^2^ flask. Lentiviral vectors for shANXA2 and shArp2 were purchased from Millipore Sigma (TRCN0000056145 and TRCN0000113861, respectively). To enhance packaging of all constructs, they were incubated for 20 minutes in polyethylenimine (3:1 ratio to DNA used) in lactate-buffered saline (pH 4.0). After 72 hrs, the lentivirus was collected and concentrated using Lenti-X^TM^ concentrator per manufacturer instructions (Clontech). Following concentration, lentivirus was suspended in M199 at 1/20^th^ of the original volume and stored at −80 ⁰C.

### Lentiviral Transduction

HUVEC were transduced with lentiviral shRNA and polybrene prior to being seeded onto gelatin-coated flasks at 50% confluency, as previously described.^34^ After five hours, growth medium was replaced and cells were incubated for four days.^41^ Knock-down was confirmed via Western blot.

### Western blotting

Cell lysates were collected in preheated Laemmli sample buffer with 2% β-mercaptoethanol. After heating for five minutes at 95°C, samples were briefly vortexed and spun down before separating via SDS-PAGE. Polyacrylamide concentration varied from 8 – 12% depending on the molecular weight of target proteins. Electrophoresis was performed at 150V, 400 amp, for one hour. Samples were then transferred onto nitrocellulose membrane at 140V, 400 amp, for 90 minutes. After transfer, blots were trimmed and placed in Western blocking solution (5% milk, Tris-buffered saline with 0.1% Tween 20) for 30 minutes, then primary antibodies were added for three hours at room temperature or overnight at 4°C. Blots were rinsed three times for five minutes in Tris-buffered saline with 0.1% Tween 20, then put in fresh Western blocking solution with secondary antibodies tagged with horseradish protein (HRP) for one hour at room temperature. Blots were washed again three times for five minutes, then HRP tags were activated with enhanced chemiluminescence solution (Millipore). Signal was detected and imaged via ChemiDoc™ MP Imaging System (Bio-Rad).

### Collagen I Preparation

Collagen matrices were prepared using collagen type I isolated from rat tail tendon as previously described.^40^ After collection, suspended collagen was lyophilized, resuspended in 0.1% acetic acid at 7.1 mg/ml stock concentration and stored at 4 ⁰C.

### Seeding Coverslips

Glass coverslips were placed in 24-well plates and coated with 50 µl collagen I (20 µg/ml) for one hour. HUVEC were then seeded (25,000 cells/50 µl M199) and allowed to adhere for 15 minutes in 5% CO_2_ at 37°C. After 15 minutes, cells on coverslips were gently mixed with a pipette for even distribution and allowed to adhere an additional 15 minutes. The cells were then given 500 µl of a 1:1 mix of M199 and growth medium, and allowed to grow overnight. The next day, cells were serum starved in M199 with Reduced Serum II^42^ for at least one hour and treated according to experimental design. After treatment, the coverslips were fixed in 4% paraformaldehyde in PBS for 20 minutes. Following two 15-minute washes in Tris-glycine, cells were permeabilized in 0.5% Triton X-100 in PBS for 20 minutes, then blocked with 1% bovine serum albumin, 1% goat serum, 0.2% sodium azide, and 0.1% Triton X-100 in PBS overnight at 4°C.

### Invasion Assay

To observe the ability of HUVEC to invade, 2.5 mg/ml collagen matrices with 1μM S1P were prepared in 96-well A/2 plates or customized silicone plates as previously described.^40, 43^ After 30 minutes of equilibration at 37⁰C in 5% CO_2_, HUVEC were seeded at 30,000 cells per 100 µl of M199 containing 50 μg/ml ascorbic acid, 40 ng/ml VEGF, 40 ng/ml bFGF, and Reduced Serum II, as described.^40, 42^ Cells were allowed to invade for the allotted time, then fixed in 3% glutaraldehyde or 4% paraformaldehyde in PBS. If fixed with glutaraldehyde, collagen matrices were rinsed with water twice prior to toluidine blue staining. Samples fixed with paraformaldehyde were rinsed with Tris-glycine twice for 15 minutes, permeabilized with 0.5% Triton X-100 for 30 minutes, and incubated in immunofluorescence blocking solution (1% bovine serum albumin, 1% goat serum, 0.2% sodium azide, and 0.1% Triton X-100) overnight and stored at 4°C.

### 2D Immunofluorescence

Coverslips were probed with primary antibodies (final concentration 1 – 4 µg/ml) in immunofluorescence blocking solution (see above) at 4⁰C overnight, then washed for one hour in 0.1% Triton X-100 in PBS, changing the wash buffer every 20 minutes. Secondary AlexaFluor antibodies (488 or 594) of the appropriate species were then added at room temperature for a one hour incubation. After a final one hour wash, DAPI (100 μM) was added for 10 minutes, then coverslips were mounted onto a slide with Fluoro Gel (Electron Microscopy Sciences) for confocal imaging.

### 3D Immunofluorescence

After incubation in immunofluorescence blocking solution, collagen matrices were placed with primary antibodies (final concentration 1 – 4 μg/ml) in fresh immunofluorescence blocking solution for three hours at room temperature. Samples were then washed in 0.1% Triton X-100 in PBS for two hours, changing the wash buffer every 20 minutes. Collagen matrices were incubated with secondary antibodies (AlexaFluor 488, 594, and/or 647; 1:600 in immunofluorescence blocking solution) for one hour at room temperature. They were then washed in 0.1% Triton X-100 in PBS for two hours, changing the buffer every 20 minutes. Samples underwent an additional wash overnight in fresh 0.1% Triton X-100 in PBS before labeling with DAPI and imagining via confocal microscope.

### Antibodies

ANXA2, Santa Cruz sc28385, http://antibodyregistry.org/AB_626677; Arp2, Abcam ab47654, http://antibodyregistry.org/AB_1139848; α-actinin 1, Santa Cruz sc17829, http://antibodyregistry.org/AB_626633; α-catenin, Sigma C2081, http://antibodyregistry.org/AB_476830; β2M, Santa Cruz sc46697, http://antibodyregistry.org/AB_626749; β-catenin, Sigma C7082, http://antibodyregistry.org/AB_258995; FAK, Cell Signaling 3285, http://antibodyregistry.org/AB_2269034; p120, Santa Cruz sc23873, http://antibodyregistry.org/AB_2086394; paxillin, Santa Cruz sc365059, http://antibodyregistry.org/AB_10844193; Phalloidin-FITC, Life Technologies A12379; VE-cadherin, Santa Cruz sc9989, http://antibodyregistry.org/AB_2077957; VE-cadherin, Abcam ab33168, http://antibodyregistry.org/AB_870662; vinculin, Sigma V9131, http://antibodyregistry.org/AB_477629; zyxin, Sigma ABC1387-25UL, https://antibodyregistry.org/AB_2905521.

### Filipin III labeling

Filipin III stocks were made in ethanol and stored in aliquots at −80 °C according to kit instructions (Abcam, ab133116). When ready for use, a working solution was made at 1:100 in PBS. Coverslips were placed cell-side down on parafilm with 20 µl of the working solution for one hour at room temperature, followed by two 10 minute washes in PBS. Coverslips were immediately transferred to a PBS-filled well in a glass-bottom plate and imaged.

### Imaging

Phase contrast images of toluidine blue stained sprouts were obtained using an Olympus CKX41 microscope, Olympus QColor3 camera, and QCapture Pro acquisition software (2048 x 1536 resolution). Density counts were captured with a 10x objective, while invasion distance was obtained from sliced matrices imaged at 20x. Confocal images of fluorescently labelled samples were obtained using a Nikon TE 2000 confocal microscope, Nikon A1 camera, and NIS-Elements acquisition software. All confocal images were obtained at 1024 resolution, with 1.4 numerical aperture. Whole field images of cell monolayers were obtained using a 60x oil immersion objective, with the exception of images for FiberFit Software™ analysis which used a 100x oil immersion objective. For imaging cells invading into collagen matrices, a 40x water immersion objective was used. To image filipin III labeled cells, monolayers were imaged on a Nikon Eclipse TE2000-U microscope with 60x oil immersion objective, using an ANDOR Zyla 4 sCMOS camera. Excitation was provided by an X-cite 120 Fluorescence Illumination System, which was managed by a Prior ProScan II controller and Nikon Elements.

### Image Analysis

With the Fiji package in ImageJ, 16-bit Nikon image files (*.nd2) were opened using the Bio-Formats plugin, then maximum intensity projections (MIPs) or 3D projections were created from the Z-stack. For colocalization measurements, MIPs were split into individual color channels, then analyzed using the Coloc2 plugin. If a particular antibody signal had high background or non-specific nuclear labeling, regions of interest were defined prior to analysis, typically using VE-cadherin signal to make a junctional mask. For threshold analysis, MIPs were made and an equal threshold applied to all images for individual experiments. Using the “Analyze Particles” tool, a map was generated of regions that passed the threshold and collected area measurements of individual objects. Image enhancement for publication was applied linearly and equally to all images within a given figure, *but did not alter actual values used for analysis*.

### Actin Fiber Analysis

FiberFit Software™, created by the Northwest Tissue Mechanics Laboratory at Boise State University, was used to measure actin fiber organization.^44^ Images were cropped into square ROIs as required by the software, which also limited analysis to junctional regions. Once converted to 8-bit color depth, images were uploaded into FiberFit Software™ and analyzed for fiber distribution.

### Super Resolution Radial Fluctuation Analysis (SRRF)

To optimize images of actin fibers prior to analysis by FiberFit Software™, we utilized SRRF. This method, established by the Henriques lab, allowed us to overcome limitations of standard confocal imaging by accounting for fluorophore movement, blinking, and the refraction limit which blur the image.^45^ By capturing 100 confocal images of the same field and Z plane, SRRF generates a sharpened representative image of fiber signal with increased resolution and contrast.

### Sample Preparation for Lipidomics

One T75 of HUVEC was transduced with shβ2M or shANXA2 lentivirus seeded onto a T175, and cultured for four days until confluent. After serum starving for one hour, cells were activated with 1 µM S1P for an additional hour. Media was removed and cells were rinsed with HEPES buffer before being collected with trypsin, which was neutralized in equal parts FBS. Cells were diluted with M199, spun down at 300 x g for 5 minutes at 4 °C, suspended in 1 ml of new M199, transferred to a 1.7 ml tube, and spun down again. The media was removed and cell pellets of approximately 12 million cells were frozen using a methanol/dry ice slurry and stored at −80 °C. Lipidomic analyses were performed by Creative Proteomics (Shirley, NY, USA). Cell pellets were thawed and suspended in 1.5 ml Chloroform/MeOH at 2:1 (v/v), vortexed for 1 minute, and sonicated at 4 °C for 30 minutes. Samples were centrifuged at 3,000 rpm at 4 °C for 10 minutes. The lower phase was transferred to a new tube and dried under nitrogen. Dried extract was suspended in 200 µl isopropyl alcohol/MeOH at 1:1 (v/v). 10 µL LPC (12:0) was added as an internal standard. After a final centrifuge at 12,000 rpm at 4 °C for 10 minutes, the supernatant was transferred for LC-MS analysis.

### LC-MS Analysis

Lipid separation was accomplished using an Ultimate 3000 LC combined with Q Exactive MS (Thermo) and screened via ESI-MS. The LC system utilized ACQUITY LPC BEH C_18_ (100 x 2.1mm x 1.7µm) with Ultimate 3000 LC. Mobile phase comprised of solvent A (60% ACN + 40% H_2_O + 10 mM HCOONH_4_) and solvent B (10% ACN + 90% isopropyl alcohol + 10 mM HCOONH_4_) with a gradient elution (0 - 10.5 min, 30% - 100% B; 10.5 min - 12.5 min, 100% B; 12.5 - 12.51 min, 100% - 30% B; 12.51 - 16.0 min, 30% B). Flow rate was 0.3 mL * min^-1^. Column temperature was maintained at 40 °C with sample manager set to 4 °C. Mass spectrometry parameters for ESI+ and ESI- mode were as follows:

ESI+: Heater Temp 300 °C; Sheath Gas Flow rate, 45 arb; Aux Gas Flow Rate, 15 arb; Sweep Gas Flow Rate, 1 arb; spray voltage, 3.0KV; Capillary Temp, 350 °C; S-Lens RF Level, 30%.

ESI-: Heater Temp 300 °C, Sheath Gas Flow rate, 45arb; Aux Gas Flow Rate, 15 arb; Sweep Gas Flow Rate, 1 arb; spray voltage, 3.2KV; Capillary Temp, 350 °C; S-Lens RF Level, 60%.

### Multivariate Statistical Analysis

Peak data was merged and imported into SIMCA-P software for analysis. Principal Component Analysis (PCA) was used followed by Partial Least Squares-Discriminant Analysis (PLS-DA) and Orthogonal Projections to Latent Structures Discriminant Analysis (OPLS-DA) to analyze observations acquired in both ion modes. Significantly different values between groups were then filtered out using VIP values > 1.5.

### Single Variable Statistical Analysis and Clustering

Lipids with a VIP > 1.5, fold change > 2.0, and p-value < 0.05 were selected as significant values for hierarchical clustering analysis. Mean values of lipid contents from biological replicates were used to calculate lipid ratios between shβ2M and shANXA2 samples. Data was adjusted to log scale and median centered ratios were normalized. Hierarchical Clustering Analysis (HCA) was performed using Cluster 3.0 (Stanford University) and visualized using Treeview (Stanford University). Lipid ratios from two independent experiments were used for HCA, where color intensity correlates with increase and decrease relative to mean ratios.

### Pathway Analysis

A correlation network diagram using the KEGG databases was constructed (Creative Proteomics), enabling the categorization of lipids to identify potential pathways impacted by loss of ANXA2.

### Figure Creation

All figures were made via a combination of ImageJ, Microsoft PowerPoint, and GraphPad Prism.

### Lipid supplementation

DPPC and DOPC were purchased from Avanti Polar Lipids (Alabaster, AL, USA) (cat. #850355 and #850375). Lipids were dried with a nitrogen stream and suspended as a 100 mM stock in 5 uM BSA and water. Stocks were sonicated in a water bath at 37 °C for 90 minutes and stored at −20 °C. To avoid possible effects from lipid oxidation, 100 mM stocks were used within one week.

### Statistics

Within a given experiment, at least three fields per treatment were obtained and measured according to experiment design. When comparing one treatment against a control, statistical significance was determined using Student’s t-test (two-tailed) where p < 0.05 was considered significant under the assumption of equal variance. When comparing multiple treatments against a control, ANOVA with Dunnett’s multiple comparisons test was performed where p < 0.05 was considered significant. All experiments were independently repeated at least three times and show similar results.

### Supplemental Materials

Additional images provide clarity regarding immunofluorescence experiments, the utility of SRRF improving the accuracy of FiberFit Software™, and information regarding lipidomic analyses.

## Results

### ANXA2 knock-down results in excess sprout initiation, but failed sprout maturation

As ANXA2 stabilizes adherens junctions and interacts with F-actin, we wanted to better visualize why ANXA2 is fundamental for endothelial sprouting in collagen matrices. We therefore performed a time series of invasion and observed localization of VE-cadherin and F-actin at 1, 5, 10, and 20 hours. After 1 hour of activation, shANXA2 cells extended twice as many initial F-actin processes into the collagen matrix compared to shβ2M controls (Fig. 1A, D). However, as time progressed, shANXA2 cells were unable to form mature sprouts, compared to control cells which formed multicellular structures. These data show that ANXA2 functions to temper the initiation of sprouts. Despite excess sprout initiation with loss of ANXA2 (1 hour), there is also failed sprout maturation over time (20 hours) as determined by formation of fewer multicellular structures that possess distinct VE-cadherin positive junctions (Fig. 1B, C, E). Additionally, because the Arp2/3 complex has been established as a critical component of sprout maturation,^46^ we also visualized localization of Arp2. Loss of ANXA2 caused reduced Arp2 in sprouts between 10 – 20 hours of invasion, so we reasoned ANXA2 may regulate localization of Arp2 and organization of F-actin to stabilize junctions. Further, careful examination of junctional changes in the absence of ANXA2 reveal wider, destabilized junctions, which is consistent with junctional destabilization caused by loss of ANXA2 (Fig. S1).

**Figure 1.**
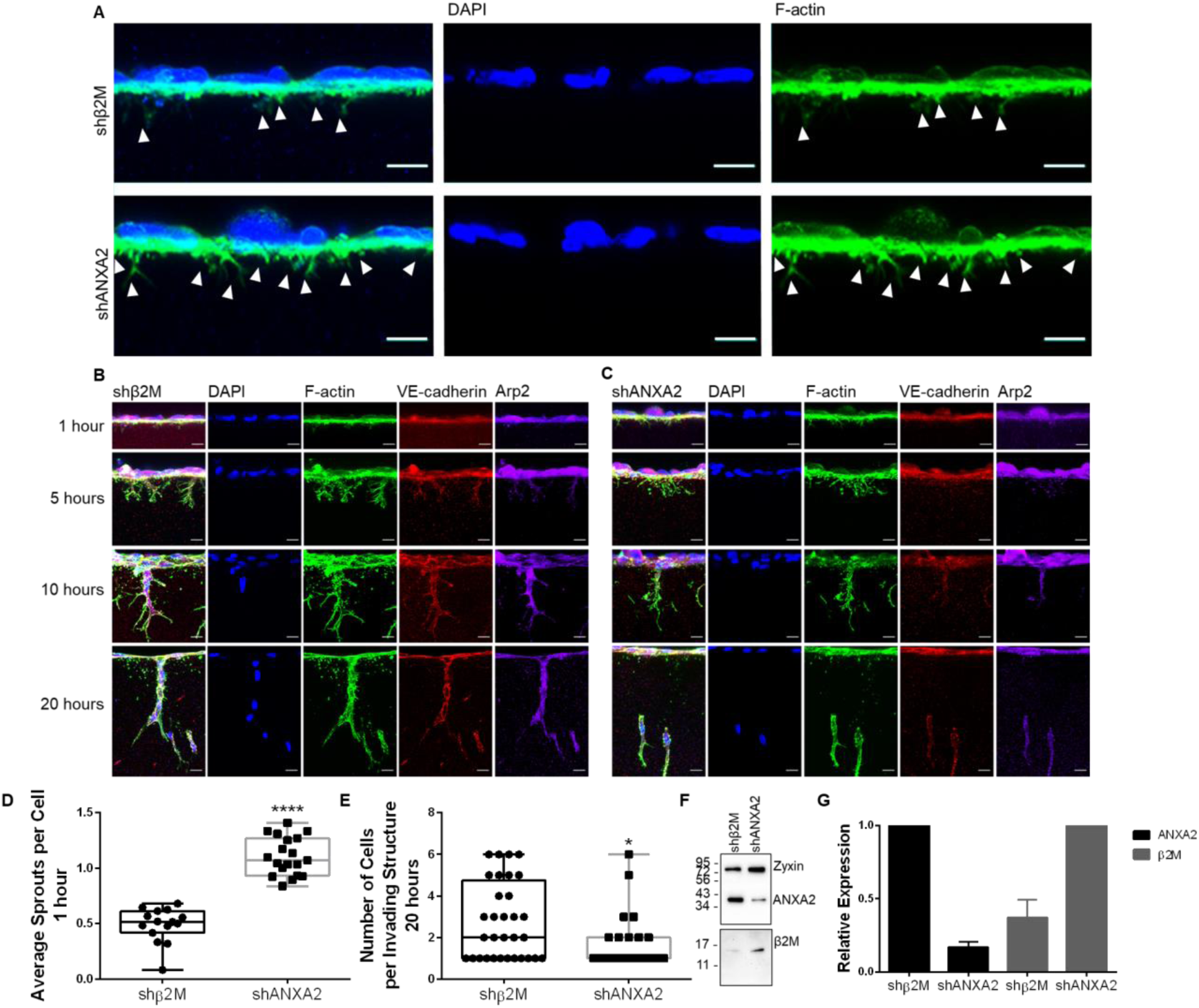
ANXA2 knock-down results in excess sprout initiation, but failed maturation. **A)** Representative side views of shβ2M and shANXA2 HUVEC on collagen I matrix after 1 hour with S1P + growth factors (GFs). Arrowheads denote F-actin (green) processes entering the matrix. Scale bar: 10 μm. **B)** Representative side views of shβ2M HUVEC on collagen I matrix after 1, 5, 10, or 20 hours of invasion with S1P + GFs. Stained for nuclei with DAPI (blue), F-actin (green), VE-cadherin (red), and Arp2 (purple). Scale bar: 10 μm. **C)** Representative side views of shANXA2 HUVEC on collagen I matrix after 1, 5, 10, or 20 hours of invasion with S1P + GFs. Scale bar: 10 μm. **D)** Average number of initiating sprouts per cell at 1 hour for fields of equal cell density. Student’s t-test, ****, p < 0.0001. n = 15, 40x fields per treatment. **E)** Quantified average cell count per invading structure after 20 hours. Student’s t-test, *, p = 0.0104. n = 25 sprout structures. shANXA2 cells fail to incorporate into multicellular structures that possess stabilized, linear junctions as labeled by VE-cadherin. **F)** Western blot of ANXA2 and β2M, with Zyxin (a focal adhesion protein) as a loading control. **G)** Normalized signal of ANXA2 and β2M from western blots of three independent experiments as confirmation of knock-down. Error bar denotes standard deviation. All experiments repeated three times with representative data shown here.

### Endothelial sprout initiation favors junctions

To understand why junctional destabilization associated with loss of ANXA2 would result in excess sprouting initiation, we imaged VE-cadherin and F-actin in wild-type cells during the early stages of sprout initiation (Fig. 2A, B). Examination of initiating sprouts relative to junctions revealed an average of 89% of sprouts initiating from adherens junctions (Fig. 2C), indicating junctions are strongly favored to support sprout development. Interestingly, the junctions where sprouts initiated (arrowheads, Fig. 2A, Fig. S2) often appeared destabilized, reminiscent of junctions in shANXA2 cells after one hour of S1P treatment (Fig. S1). This supports the concept that adherens junctions must locally destabilize prior to sprout formation, taking on a reticular morphology as junctional complexes remodel to initiate invasion.

**Figure 2.**
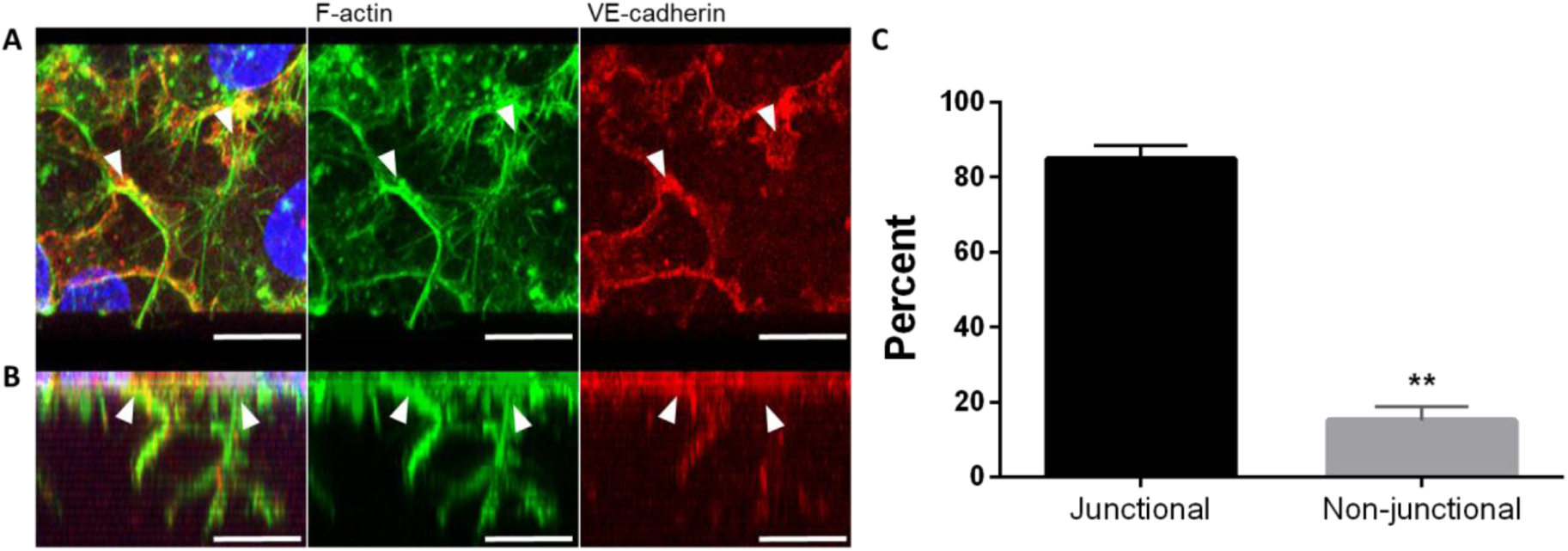
Endothelial sprout initiation favors junctions. **A)** Top view of HUVEC seeded on collagen matrices, activated with S1P for one hour, then VEGF and bFGF for an additional hour. Cells were labeled for F-actin (green), VE-cadherin (red). Nuclei stained with DAPI (blue). **B)** Side view of 3D reconstruction from the Z-stack in A. White arrowheads denote origin points of sprouts. Scale bars: 10 μm. **C)** Bar graph of average sprout origins (percent) of three independent experiments, +/- standard deviation. Student’s t-test: **, p = .0036, n = 375 sprouts total.

### ANXA2 knock-down causes disorder of actin fibers at junctions during S1P signaling

Because ANXA2 is known to assist with actin bundling,^47^ and bundled actin is associated with stable junctions,^48^ we considered that loss of ANXA2 may result in alterations of F-actin organization which would further contribute to junctional destabilization. We compared F-actin organization relative to VE-cadherin positive junctions in shβ2M and shANXA2 cells after one hour of S1P activation (Fig. 3). To quantify actin organization, we utilized FiberFit Software™ to analyze the relative degree of order within established junctional regions between exactly two cells (Fig. 3B).^44^ We also used super-resolution radial fluctuation (SRRF) analysis to optimize the clarity of phalloidin signal for the purposes of increasing image contrast.^45^ This allowed the threshold application by FiberFit Software™ to detect individual fibers more accurately (Fig. S3). The analysis revealed that shANXA2 cells show decreased order of F-actin fibers (K) and decreased fit to normal curve (R^2^), indicating increased disorganization of F-actin fibers compared to the parallel cortical fibers that are characteristic of S1P signaling in shβ2M controls (Fig. 3C). We also observed a decrease in the size of focal adhesions at junctions (Fig. S4), which suggests there is reduced tensional force at the cell periphery to support junctions.^49, 50^ Altogether, this validates that shANXA2 disorganization of F-actin and destabilization of junctions favors a state of remodeling for sprout initiation.

**Figure 3.**
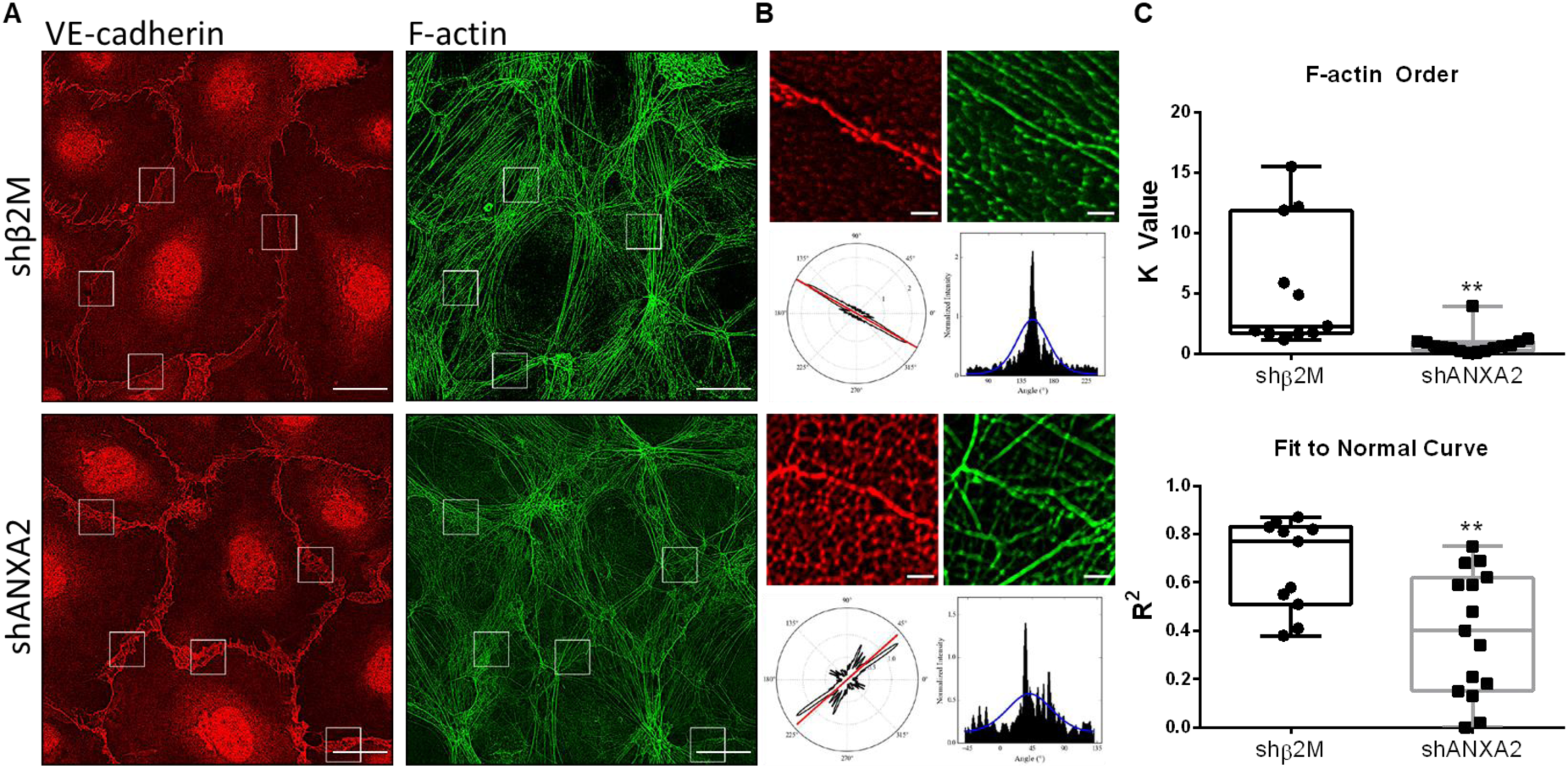
ANXA2 knock-down results in disorder of actin fibers at junctions following S1P treatment. **A)** Whole field of shβ2M and shANXA2 cells on coverslips following one hour S1P treatment, labeled for VE-cadherin and F-actin. White boxes denote ROIs chosen for analysis in FiberFit SoftwareTM. Scale bar: 10 μm. **B)** Example ROIs that were loaded into FiberFit Software™ with corresponding VE-cadherin channel as reference (only F-actin channel was analyzed). Lower left panel denotes radial graph of F-actin signal. Lower right graph is the radial graph plotted as a histogram. Blue line denotes Gaussian curve. Scale bar: 1 μm. **C)** Averaged results from B. Top graph shows the relative degree of order, quantified as K value. n = 3 fields, at least 12 junctions per treatment. **, p = 0.002, Student’s t-test. Bottom graph shows fit to normal curve, reflecting average fit of the blue line in B. **, p = 0.0053, Student’s t-test. All experiments independently repeated at least three times with representative data shown.

### Localization of Arp2, α-actinin 1, and vinculin is altered with ANXA2 knock-down

Since the organization of F-actin is disrupted with ANXA2 knock-down, we considered whether other actin-regulating proteins fail to localize to junctions in response to S1P. To evaluate actin nucleation/branching, bundling, and organization, we analyzed Arp2,^51, 52^ α-actinin 1,^53, 54^ and vinculin,^55^ respectively. In shANXA2 cells, we observed significantly less localization of Arp2 at junctions, as quantified by Pearson’s Coefficient (Fig. 4A, B). α-actinin 1 localized at junctions similarly to control cells, however its organization and distribution reflected the reticular pattern of VE-cadherin and disorganization of actin fibers from loss of ANXA2 (Fig. 4C, D). We also found that vinculin localization did not change relative to VE-cadherin, but its organization maintained association with disorganized junctions from loss of ANXA2 (Fig. 4E, F). It is interesting to note that loss of the Arp2/3 complex at junctions implies that the generation of new cortical fibers is reduced, as there would be reduced assembly of localized lamellipodia, resulting in reduced structural reinforcement of junctions.^56^ Altogether these findings indicate that the organization of actin fibers relative to junctions fails, which is in agreement with figure 3.

**Figure 4.**
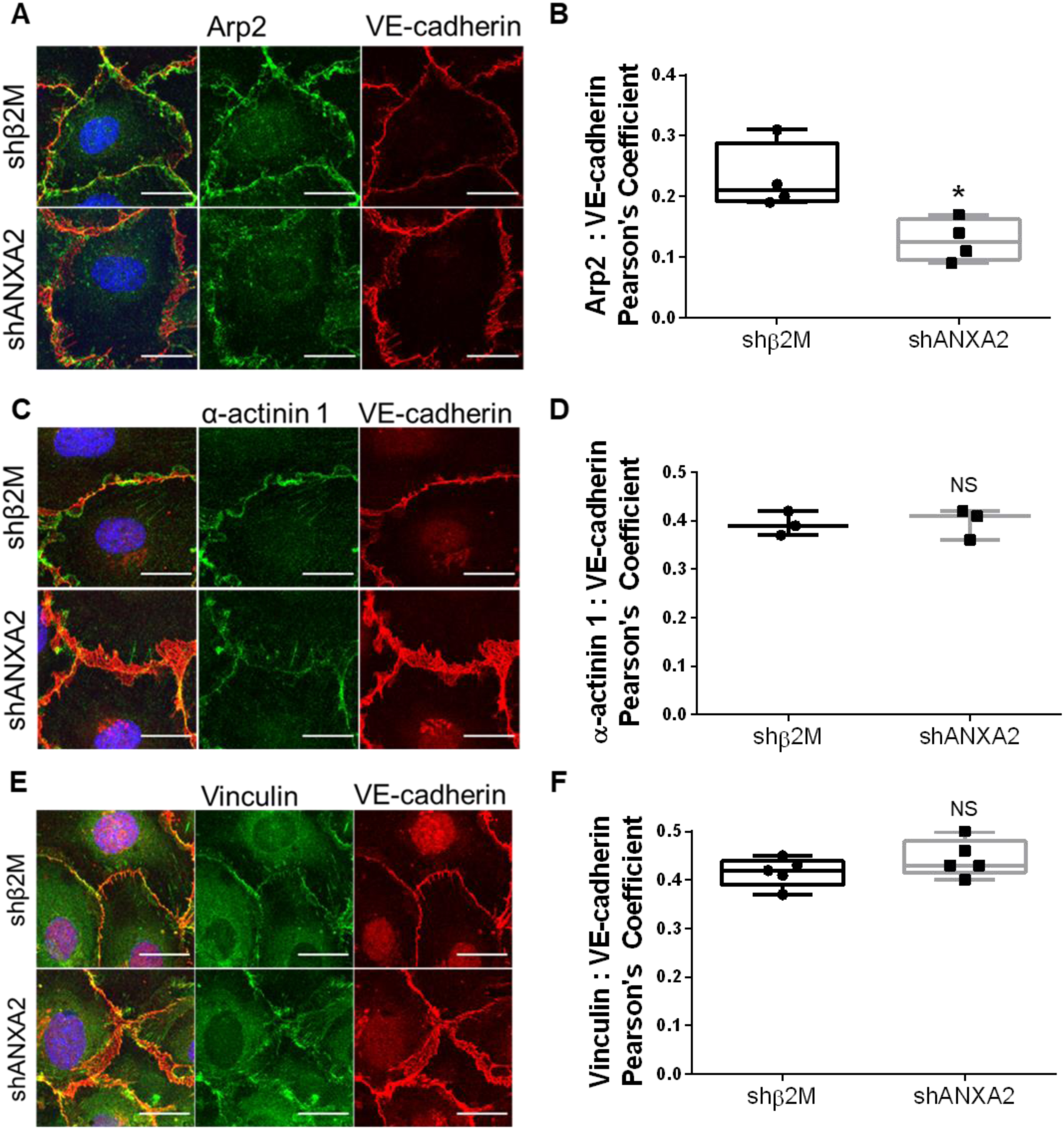
Junctional localization of Arp2, α-actinin 1, and vinculin with ANXA2 knock-down. **A)** Immunofluorescence of shβ2M and shANXA2 cells on coverslips following one hour S1P treatment. Cells were then labeled for Arp2 and VE-cadherin. **B)** Quantified colocalization between Arp2 and VE-cadherin as measured by Pearson’s Coefficient. n = 4 fields per treatment; Student’s t-test: *, p = 0.0197. **C)** Immunofluorescence of shβ2M and shANXA2 cells after S1P treatment, labeled for α-actinin 1 and VE-cadherin. **D)** Pearson’s Coefficient as quantification of colocalization between α-actinin 1 and VE-cadherin. n = 3 fields per treatment; Student’s t-test: NS = no significance. **E)** Immunofluorescence of shβ2M and shANXA2 cells after one hour S1P, labeled for Vinculin and VE-cadherin. **F)** Colocalization of Vinculin and VE-cadherin, measured by Pearson’s Coefficient. n = 5 fields per treatment; Student’s t-test: NS = no significance. Scale bars: 10 μm. All experiments repeated independently three times with representative data shown.

### Loss of Arp2 impairs sprouting maturation, but does not affect initiation, cortical F-actin order, or ANXA2 localization

To better understand the interplay between ANXA2, Arp2, and actin dynamics, we reduced Arp2 expression using a short-hairpin RNA (shArp2). After confirmation of knock-down, we performed invasion assays and found that loss of Arp2 results in impaired sprouting (Fig. 5A-F). Unlike shANXA2 cells, we did not observe excessive sprout initiation at one hour (Fig. 5G, H), however we did observe a change in formation of multicellular structures after overnight invasion. The shArp2 group contained fewer cells and less continuous VE-cadherin-positive junctions compared to the shβ2M group (Fig. 5I, J). In 2D immunofluorescence experiments, after one hour of S1P activation, we surprisingly saw no significant change in junctional morphology with loss of Arp2, nor increased disorder of cortical F-actin (Fig. 6A-D). Finally, localization of ANXA2 was not altered with loss of Arp2 (Fig. 6E, F); altogether this suggests that ANXA2 functions in a manner upstream of Arp2 organization, and Arp2 is vital for the maturation of multicellular invading structures but not sprouting initiation or cortical actin organization.

**Figure 5.**
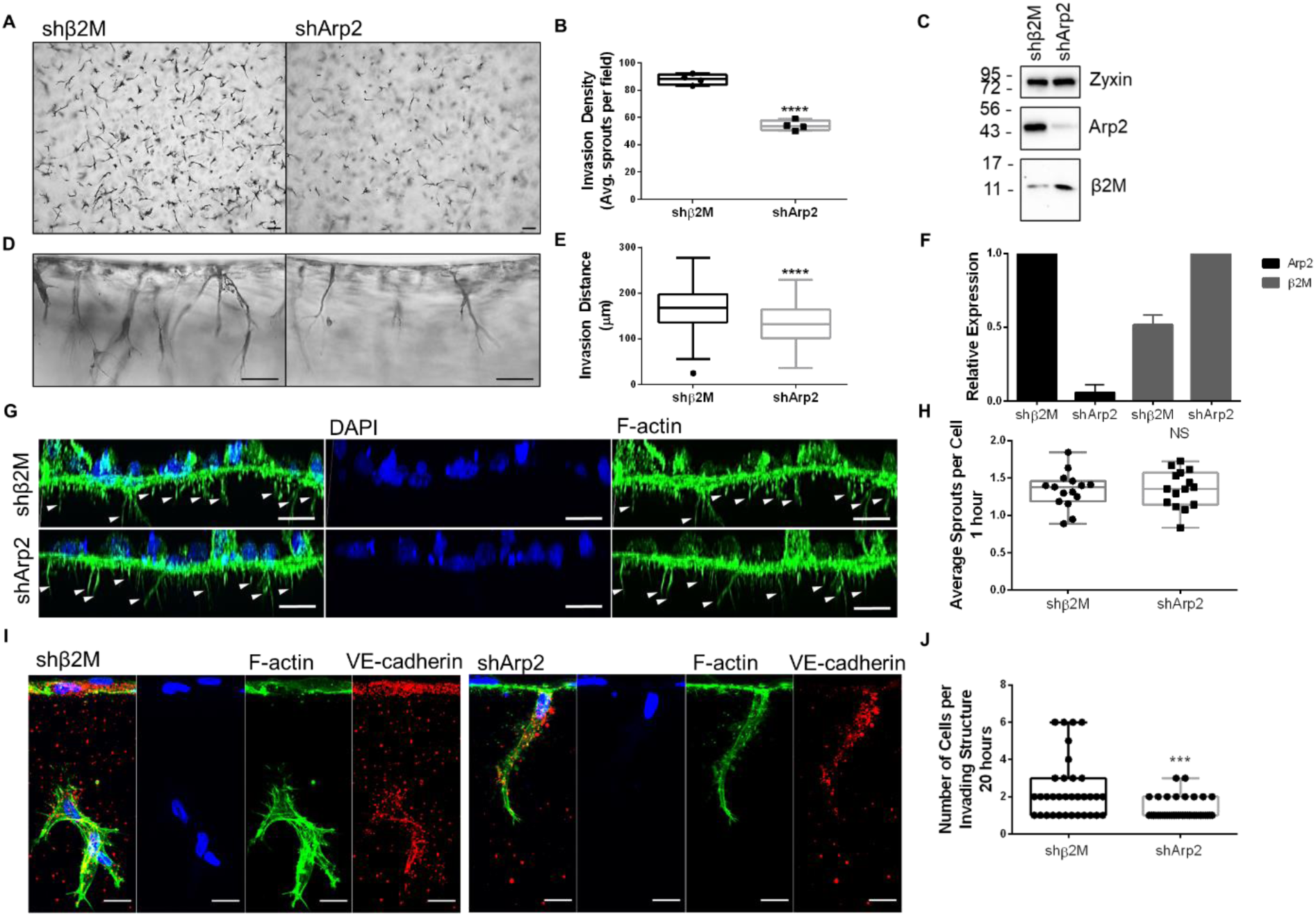
Arp2 is required for sprouting in 3D collagen matrices, but does not effect sprouting initiation. **A)** Top view of sprouts in collagen matrices after 21 hours. Scale bar: 100 μm. **B)** Quantified sprouts per field; n = four fields per group. Student’s t-test: ****, p < 0.0001. **C)** Western confirmation of Arp2 knock-down. **D)** Side view of sprouts in collagen matrices after 21 hours. Scale bar: 100 μm. **E)** Quantified invasion distance. n = 100 sprouts per group; Student’s t-test: ****, p < 0.0001. **F)** Relative expression of Arp2 and β2M from three independent experiments. **G)** 3D reconstructed Z-stack of F-actin (green) in shβ2M or shArp2 cells after 1 hour invasion. Arrowheads denote initiating processes. **H)** Average number of sprouts per cell after 1 hour. n = 15 fields per treatment. Student’s t-test: p > 0.05, NS, not significant. **I)** Immunofluorescence labeling of F-actin (green) and VE-cadherin (red) in shβ2M and shArp2 cells after overnight invasion. Scale bar = 10 µm. **J)** Average number of cells per invading structure at 20 hours. n ≥ 35 sprouts per treatment; Student’s t-test: ***, p = 0.0003.

**Figure 6.**
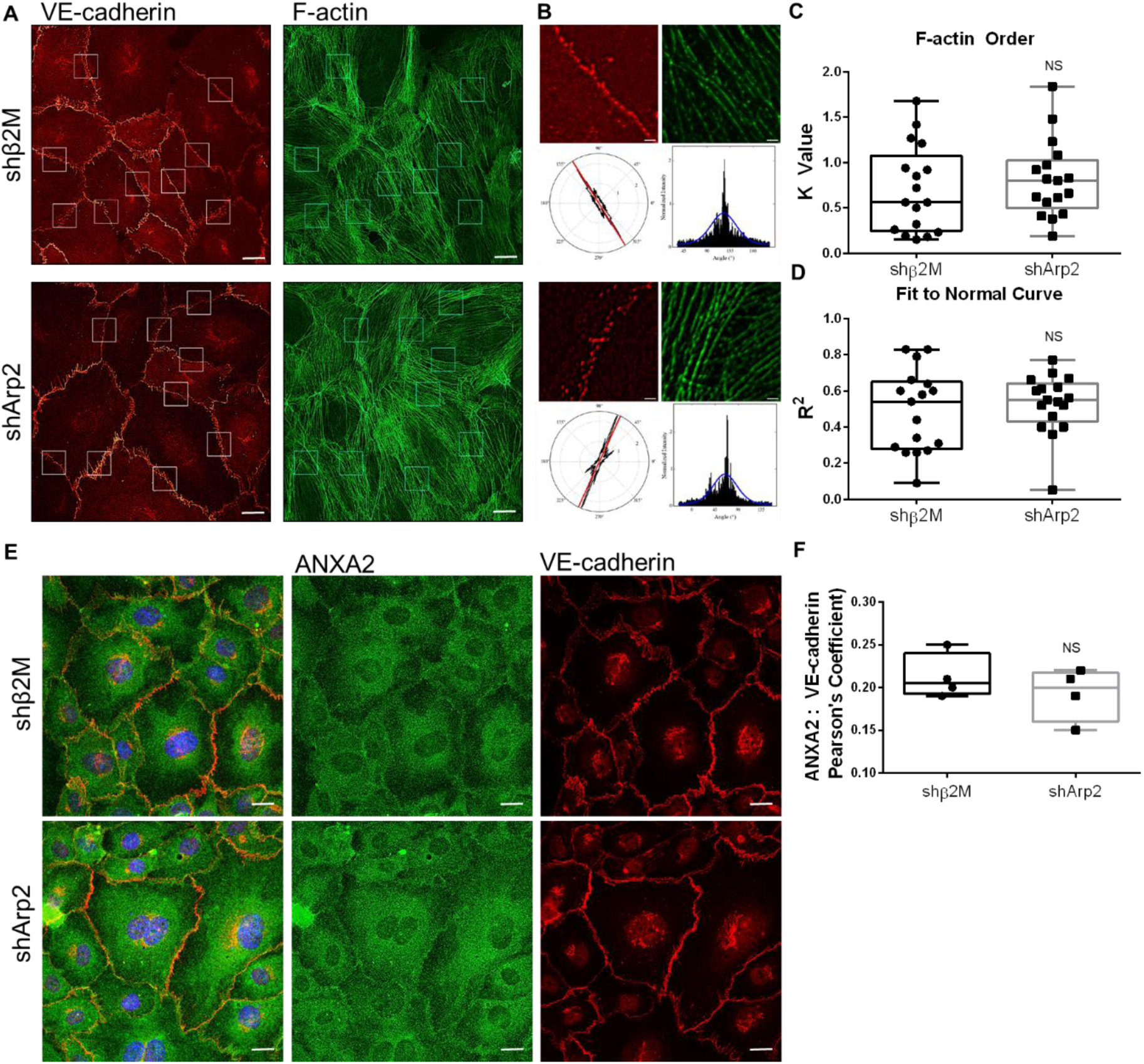
Loss of Arp2 does not impact organization of cortical F-actin or localization of ANXA2. **A)** Immunofluorescence of shβ2M and shArp2 cells on coverslips, labeled for F-actin (green) and VE-cadherin (red). Scale bar: 10 μm. **B)** Example output from FiberFit analysis. Scale bar: 1 μm. **C)** Quantified order of cortical F-actin from FiberFit. n = 17 junctional regions of interest from at 3 fields per treatment. Student’s t-test: p > 0.05, NS, not significant. **D)** Quantified Gaussian distribution of cortical F-actin from FiberFit. n = 17 junctional regions of interest from at 3 fields per treatment. Student’s t-test: p > 0.05, NS, not significant. **E)** Immunofluorescence of ANXA2 (green) and VE-cadherin (red) after one hour of S1P activation. Scale bar: 10 µm. **F)** Quantified colocalization between ANXA2 and VE-cadherin in shβ2M and shArp2 cells. Student’s t-test: p > 0.05, NS, not significant. All experiments were performed three times and representative results shown.

### The complex between filipin III and cholesterol is greatly reduced with loss of ANXA2 in intact cells, but restored with membrane disruption by permeabilization

Membrane cholesterol is known to be a critical regulator of lipid rafts involving PI(4,5)P_2_, adherens junction stability, and actin dynamics.^57, 58^ Though it has been shown *in vitro* that ANXA2 does not directly bind cholesterol^59^ and the presence of cholesterol enhances ANXA2 binding to membranes,^60, 61^ to our knowledge, an ANXA2-dependent mechanism of cholesterol function has not been identified. Initial experiments showed that the localization of filipin III, which specifically labels cholesterol, was enhanced at junctions in response to S1P (Fig. S5), so we next considered whether localization of cholesterol is dependent on ANXA2, as loss of ANXA2 results in junctional destabilization. We observed a dramatic loss of filipin III signal in shANXA2 cells, suggesting either the total level of cholesterol was reduced or cholesterol was no longer accessible to complex with filipin III (Fig. 7A, B). Disruption of the plasma membrane with TX-100 resulted in similar filipin III signal between shANXA2 and control cells (Fig. 7A, C), suggesting that cholesterol is present, however its presentation within the membrane is altered and inaccessible by filipin III. This finding was reinforced via thin-layer chromatography (TLC) on whole cell lipid extracts after one hour of S1P treatment. Using purified cholesterol as a standard, we found no significant change in total cholesterol between shANXA2 and shβ2M cells (Fig. 7D, E). This is in agreement with filipin III signal in permeabilized cells, suggesting ANXA2 maintains orientation of cholesterol within intact membranes.

**Figure 7.**
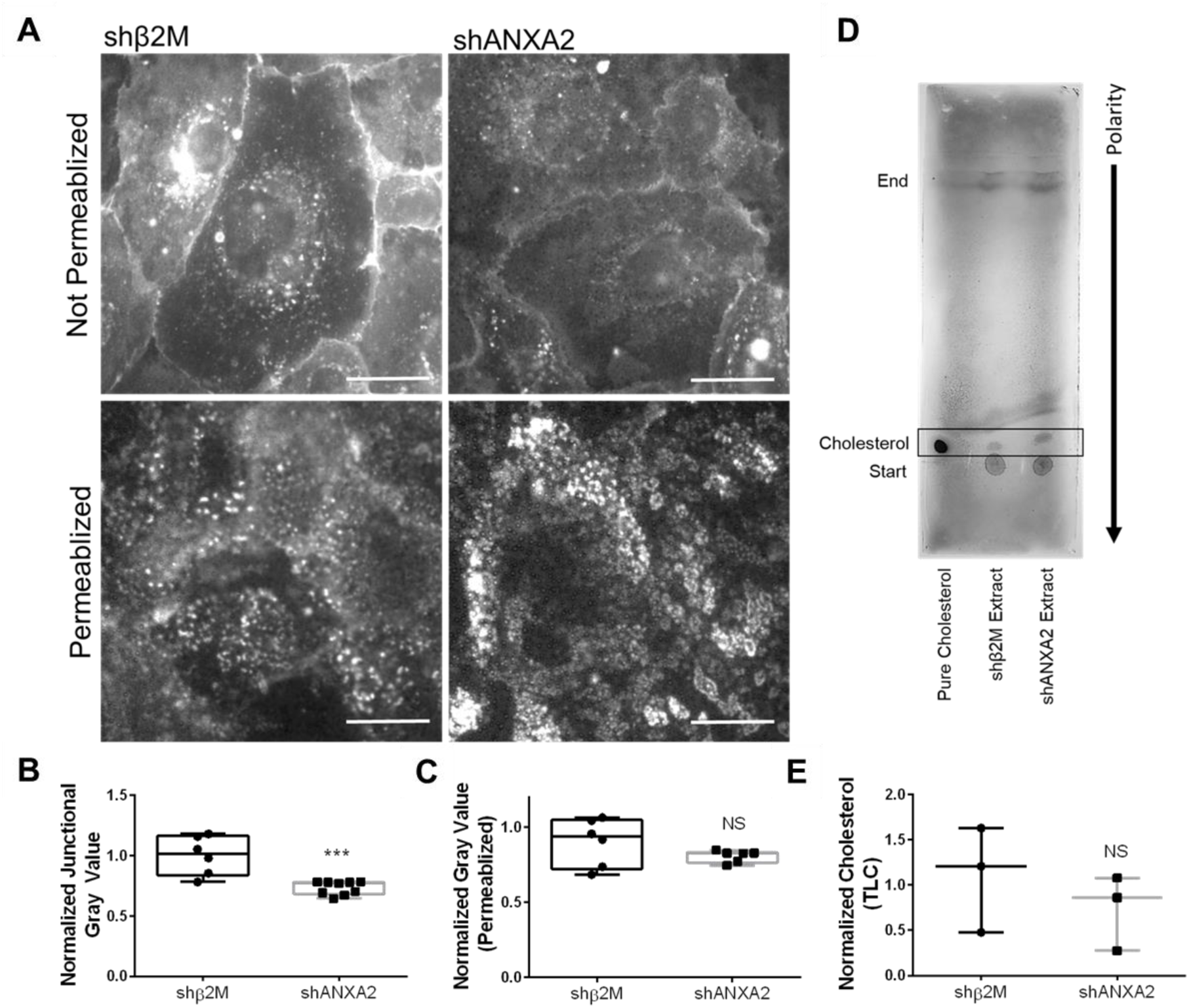
Loss of ANXA2 impairs filipin III labeling of cholesterol within intact membranes, without reducing total cholesterol. **A)** shβ2M and shANXA2 cells on coverslips after one hour of S1P treatment, stained with filipin III with or without permeabilization. **B)** Quantified junctional filipin III signal in shβ2M and shANXA2 without permeabilization. Student’s t-test, ***, p = 0.0004. n ≥ 6 fields per treatment. **C)** Quantified whole field filipin III signal in cells with permeabilization. Student’s t-test, p > 0.05, NS = not significant. n = 6 fields per treatment. **D)** Whole cell lipid extracts of shβ2M and shANXA2 cells run through thin-layer chromatography on silicate plate. **E)** Quantified cholesterol signal from B over three experiments. Student’s t-test, p > 0.05, NS = not significant.

### ANXA2 does not broadly impact lipid classes or acyl chain metabolism

Composition of the lipid bilayer dramatically impacts the behavior of membrane cholesterol,^62^ so we hypothesized that loss of ANXA2 will change membrane lipid composition. To determine this, whole cell, untargeted lipidomics was used to measure changes in lipid composition with loss of ANXA2. After one hour of S1P treatment, shβ2M and shANXA2 endothelial cells were analyzed by liquid chromatography–mass spectrometry (LC-MS). This process identified and quantified 1,894 lipid species, which were organized according to general lipid class, acyl chain length, and saturation. Overall, shβ2M and shANXA2 cells were clustered separately, however lipid classes as a whole between the two groups were not significantly changed (Fig. S6). We observed both shβ2M and shANXA2 cells contained equal parts phosphatidylethanolamine (PE) and PC at approximately 40% each, while phosphatidic acid (PA), phosphatidylinositol (PI), phosphatidylserine (PS), and phosphatidylglycerol (PG) ranged between 1 – 2% (Fig. 8A). When considering acyl chain composition, there was very little change between shβ2M and shANXA2 cells (Fig. 8B). The majority of stereospecific numbered-1 (sn-1) acyl chains were 16 and 18 carbons long and either saturated or monounsaturated. Sn-2 fatty acids were mainly 16, 18, 20, or 22 carbons long and were mostly monounsaturated, followed by saturated and polyunsaturated. There was a small, yet significant decrease detected in sn-2 acyl chains of 6 and 15 carbons, while an increase in 16 carbon sn-2 chains was detected with loss of ANXA2 (Fig. 8B). These results indicate that the majority of phospholipid synthesis/modification events are still occurring in the absence of ANXA2, and further investigation of specific lipid species is required.

**Figure 8.**
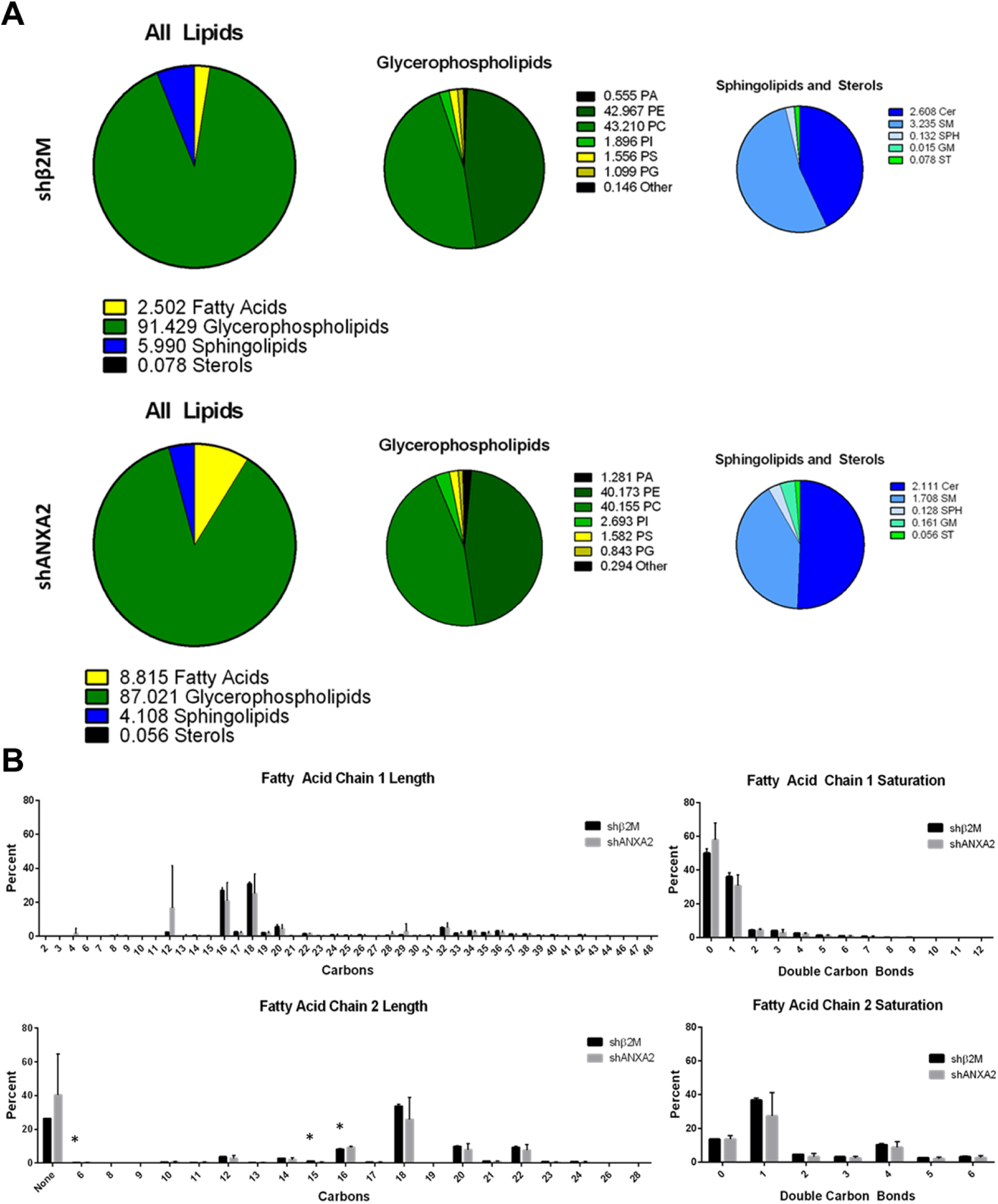
Overview of lipid profile in shANXA2 HUVEC. **A)** Breakdown of total lipid profile by lipid class, averaged between 3 cell lots. No significance detected according to Student’s t-test across the 3 cell lots. **B)** Breakdown of acyl chain length and saturation with loss of ANXA2, average between 3 cell lots. Error bars represent standard deviation. Student’s t-test: *, p = 0.021, 0.027, and 0.033 for 6, 15, and 16 carbons on sn-2, respectively. Note: Very long chains (> 28 carbons) likely include both sn-1 and sn-2 chains, which LC-MS could not identify at higher resolution under the given parameters.

### Loss of ANXA2 alters specific lipid species

Though loss of ANXA2 did not overtly impact phospholipid composition, hierarchical clustering analysis (HCA) identified significant changes in 42 lipid species with loss of ANXA2 (Fig. 9). This included reduction in several species of PC, PE, and PS, as well as minor species of sphingomyelin, ceramide, and diacylglycerol (Fig. 9A, Table 1). Of the identified lipids, only two increased with loss of ANXA2, identified as PS (42:9) and PC (16:0_16:0), also known as DPPC. Despite the analysis in figure 8 suggesting there were no significant shifts in lipid class, a correlation network was established using Kyoto Encyclopedia of Genes and Genomes (KEGG) databases. Input of the significantly altered lipid species into categorical annotations allowed for pathway enrichment analysis to highlight potentially relevant biomarker networks (Fig. S7).

**Figure 9.**
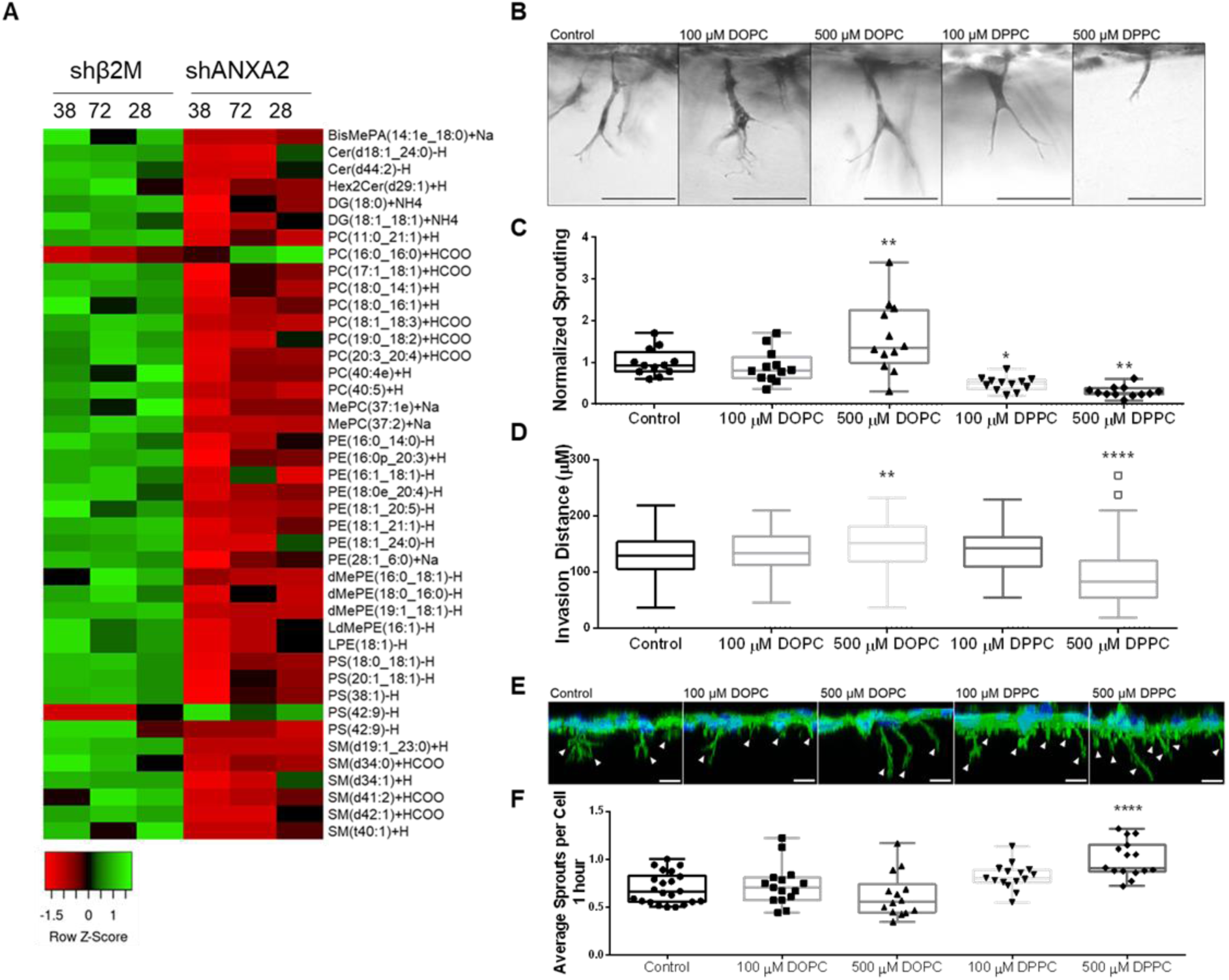
Supplementation with PC (16:0_16:0) (DPPC) impairs sprouting. **A)** Heat map of lipid species altered by loss of ANXA2. **B)** Side view of invasion after 21 hours while supplemented with DPPC or DOPC. **C)** Sprouting density normalized to the average control density. ANOVA, Dunnett’s multiple comparisons: **, p = 0.0088, *, p = 0.0241, **, p = 0.0013. n = 12 fields per treatment across 3 independent experiments. **D)** Quantified average invasion distance with lipid supplementation. ANOVA, Dunnett’s multiple comparisons: **, p = 0.0067, ****, p < 0.0001. n = 100 sprouts per treatment. **E)** 3D reconstruction from Z-stack of HUVEC after one hour of invasion. F-actin (green) reveals initiating sprouts, denoted by white arrowheads. Scale bar: 10 µm. **F)** Quantified initiating sprouts per cell after one hour. ANOVA, Dunnett’s multiple comparisons: ****, p < 0.0001. All experiments repeatedly independently at least three times with representative data here.

**Table 1.**
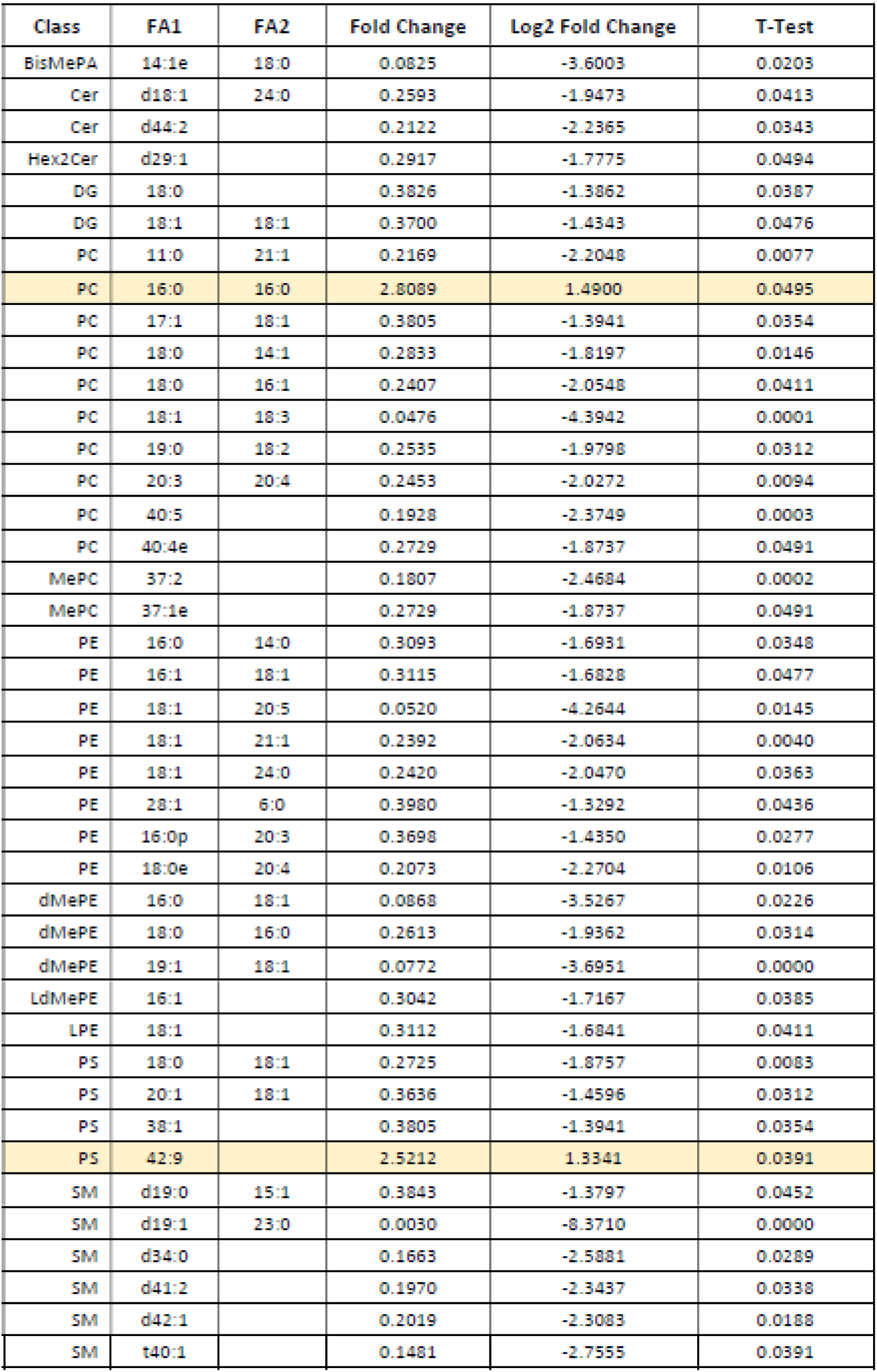
Listed fold change and significance of lipid species affected by loss of ANXA2 after 1 hour of S1P treatment. Lipids that increase with loss of ANXA2 are highlighted yellow.

### Supplementation of PC (16:0_16:0) mimics the shANXA2 phenotype

To determine if excess PC (16:0_16:0), referred from here on as DPPC, is causal or symptomatic of endothelial dysfunction, we evaluated HUVEC on coverslips and in 3D invasion assays after supplementation with DPPC. As a positive control, we used PC (18:1_18:1), referred from here on as DOPC (dioleoylphosphatidylcholine), which was one of the most abundant phosphatidylcholines detected in our system that was unaffected by loss of ANXA2. Cells were seeded onto collagen-coated coverslips or collagen matrices in the presence of these lipids to determine if there were lipid-dependent effects. Similar to ANXA2 knock-down, supplementation with DPPC decreased overnight sprouting responses, yet promoted excess sprout initiation at one hour (Fig. 9B-F). Further, DPPC supplementation caused wider junctions (Fig. 10A, B) and impaired cholesterol labeling with filipin III (Fig. 10C, D). Interestingly, elevated DOPC also slightly reduced filipin III signal, despite not having an effect on junctional morphology. DOPC did not impact the organization of F-actin, while DPPC treatment resulted in decreased order and reduced normal distribution (Fig. 10E-G). Lastly, DPPC treatment impaired the recruitment of Arp2 to junctions, as measured by colocalization with VE-cadherin (Fig. 10H, I). Together, these findings indicate that DPPC treatment mimics loss of ANXA2, inducing excessive sprout initiation, but failed sprout maturation. This is concurrent with wider adherens junctions, disrupted cortical actin organization, and failed recruitment of Arp2.

**Figure 10.**
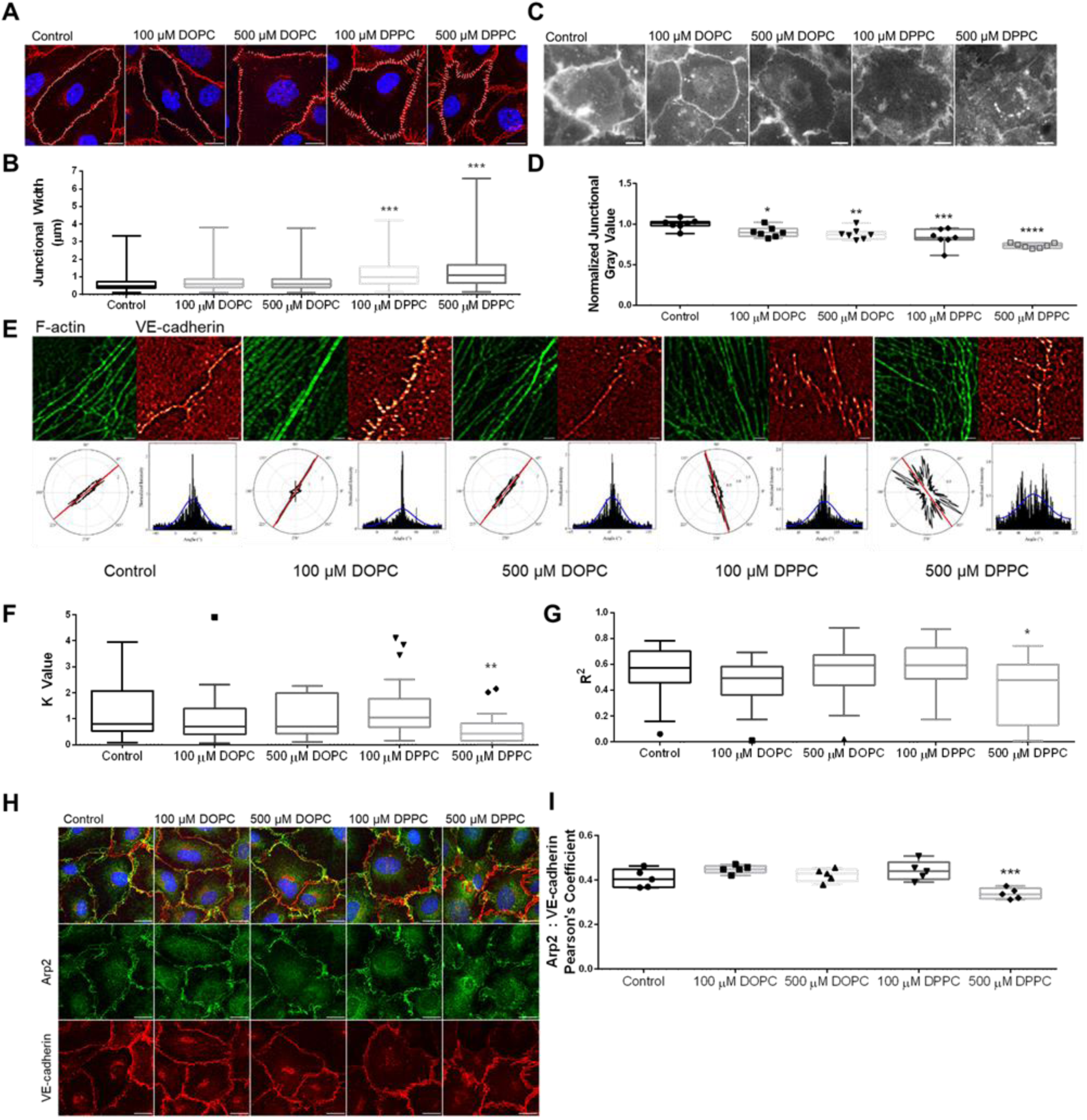
Supplementation with DPPC results in improper junctional regulation. **A)** VE-cadherin signal of wild-type cells on coverslips after overnight treatment with DPPC and DOPC, then treated with S1P for one hour. White lines denote junctional width measurement. Scale bar: 10 µm. **B)** Average junctional width from A. ANOVA, Dunnett’s multiple comparisons: ***, p = 0.0009, 0.0001 for 100 and 500 µM DPPC, respectively. n = 4 fields per treatment. **C)** Filipin III signal of HUVEC after lipid supplementation overnight and one hour S1P treatment. Scale bar: 10 µm. **D)** Average junctional gray value per field, normalized to the control. ANOVA, Dunnett’s multiple comparisons: *, p = 0.0483, **, p = 0.0093, ***, p = 0.0002, ****, p < 0.0001. n = 7 fields per treatment. **E)** FiberFit™ analysis of HUVEC seeded on coverslips overnight with DPPC or DOPC treatment, followed by one hour S1P treatment. **F)** K value of F-actin organization from FiberFit™. ANOVA, Dunnett’s multiple comparisons: **, p = 0.0048. n = 20 ROIs from 4 fields per treatment. **G)** R^2^ value of F-actin distribution from FiberFit™. ANOVA, Dunnett’s multiple comparisons: *, p = 0.0272. n = 20 ROIs from 4 fields per treatment. **H)** Arp2 localization of HUVEC after overnight lipid supplementation and one hour S1P treatment. Scale bar: 10 µm. **I)** Quantified colocalization between Arp2 and VE-cadherin using Pearson’s Coefficient. ANOVA, Dunnett’s multiple comparisons: ***, p = 0.0003. n = 5 fields per treatment. All experiments repeatedly independently at least three times with representative data shown.

## Discussion

For a cell to carry out complex morphological changes, such as sprout formation within an extracellular matrix, coordination between the membrane and protein complexes is required. Dramatic shifts in cell structure inevitably result in regions of enhanced stress, which require targeted reinforcement to overcome energy requirements and remain stable.^63^ The results shown here highlight the importance of the adaptor protein, ANXA2, in organizing these complexes. We found that ANXA2 facilitates organization of actin fibers at junctions, likely through regulation of membrane composition. When cells lacking ANXA2 are activated with S1P, there is accumulation of DPPC, which causes the actin cytoskeleton to become disorganized and adherens junctions to become destabilized. While destabilized junctions are a favored site of sprout initiation, the hyper-initiation seen with ANXA2 knock-down ultimately results in failed sprout maturation, as cells cannot assemble into multicellular structures. These results illustrate that angiogenesis is a balance between opposing dynamic forces on the junctional barrier that requires careful coordination at the membrane.

We report for the first time that ANXA2 is upstream of Arp2 recruitment to junctions, and this has particularly interesting implications for cell invasion in 3D. In cancer cells, the Arp2/3 complex is a critical component of invadopodia, the structures that assemble to promote cell invasion and degradation of the extracellular matrix.^64, 65^ In addition to the Arp2/3 complex, α-actinin and vinculin have been shown to play critical roles in invadopodia function.^66–69^ While invadopodia is a term generally reserved for cancer cells, the overlap between proteins localized to invadopodia and these responsible for endothelial cell invasion is undeniable. It is also striking that in our model, S1P selectively recruits these proteins to endothelial junctions. Junctional organization of ANXA2, Arp2, and F-actin allows these assembled protein complexes to simultaneously stabilize junctions and recruit the cellular machinery needed to invade and extend sprouts into the matrix.

Recently, invasive structures in endothelial cells were characterized and termed dactylopodia, based on similar appearance to structures in amoeboid cells.^46^ A defining attribute of the dactylopodia structure is recruitment of the Arp2/3 complex, which enables expansion of thin filopodia into a trunk that advances angiogenic development. As seen in figure 1, control cells show enhanced Arp2 signal in sprouts by 10 hours of invasion, while shANXA2 cells do not. If the Arp2/3 complex is not recruited appropriately (either to junctions or sprouts extending from junctions), then it follows that actin cytoskeletal dynamics and polymerization would be affected, resulting in a less stable structure. To our knowledge, this is the first report showing ANXA2 is necessary for proper localization of Arp2 to endothelial adherens junctions. It has been shown that ANXA2 works in cooperation with the Arp2/3 complex during F-actin assembly, endosome biogenesis, and autophagosme formation,^70–72^ however dependence of the Arp2/3 complex on ANXA2 had not been fully established.

While studies on ANXA2 have consistently implicated dynamics of F-actin and membranes, understanding the exact mechanisms remains a challenge due to limits in assaying specific lipid species. Advances in lipidomics have made the field much more accessible for systematic study across disciplines. This could allow for the identification of cell-type specific shifts in lipid profiles under a variety of conditions, which would reveal new targets for identification and treatment of disease. In this project, we provide evidence that ANXA2 regulates specific lipids, particularly DPPC, which likely impacts positioning of cholesterol and membrane properties at junctions. While it has been shown *in vitro* that ANXA2 does not directly bind cholesterol,^59^ we find loss of ANXA2 reduces the ability of filipin III to complex with cholesterol. Membrane-specific reduction of filipin labeling of cholesterol has been reported in vesicle formation,^73^ and is argued to be dependent on proper presentation of cholesterol within the membrane.

The misalignment of cholesterol within junctional membranes could be explained by changes in the 42 individual lipids identified by lipidomics in shANXA2 cells, notably DPPC, which likely induces a shift in lipid segregation. If lipid domains at the plasma membrane are not organized appropriately, transmembrane complexes may lose stability, resulting in destabilized adherens junctions. This is mimicked by the exogenous addition of DPPC, where supplementation in wild-type HUVEC resulted in wider adherens junctions and impaired sprouting angiogenesis similar to shANXA2 cells. Interestingly, DPPC is the major component of lung surfactant, which significantly alters the properties of the alveolar surface for proper lung function.^74^ This may be relevant to our observation of the effect of DPPC on endothelial adherens junctions, where in the event of damage to the lung epithelium, the endothelium would be exposed to leaking lung surfactant, opening junctions to enable recruitment of immune cells. Similarly, as DPPC has been shown to be increased in parts of the brain in an Alzheimer’s disease mouse model,^36^ its accumulation may be causal of destabilization of the blood brain barrier common with neurodegenerative pathologies. Interestingly, we were able to identify correlations between pathological conditions, expression of DPPC, and expression of ANXA2 in a variety of reports (Table 2). Both increased DPPC and decreased ANXA2 have been associated with esophageal cancer,^75, 76^ osteosarcoma,^77, 78^ and non-malignance of salivary tumors.^79, 80^ Also interesting is a possible association of DPPC and ANXA2 in both lupus and preeclampsia, where increased DPPC can accumulate to form “active hydrophobic spots,” possibly in conjunction with antiphospholipid syndrome and development of autoantibodies against ANXA2.^81–85^ The connection between ANXA2 and DPPC identified here presents exciting new possibilities for management of disease conditions. As excess DPPC can induce junctional defects in endothelial cells, it will be interesting to determine if that association persists in the conditions listed in Table 2.

**Table 2.** List of disease conditions associated with expression of ANXA2 and PC (16:0_16:0).

| **Disease** | **ANXA2** | **PC (16:0_16:0)** | **Citation** |
| --- | --- | --- | --- |
| Esophageal Cancer | Decreased | Increased | [75, 76] |
| Lupus | Increased<br>(Autoantibodies) | Increased<br>(Active Hydrophobic Spots) | [81, 82] |
| Non-malignant Salivary Tumor | Decreased | Increased | [79, 80] |
| Osteosarcoma | Decreased | Increased | [77, 78] |
| Preeclampsia | Decreased | Possibly Increased<br>(Identified as PC (32:0)) | [84, 85] |

The status of the junctional barrier remains a critical determinant of successful angiogenesis. As shown here, junctional destabilization facilitates sprout initiation, while maintenance of stabilized junctions is necessary for sprout maturation. This is likely modulated through proper phospholipid composition, which provides a foundation from which to organize multicellular sprouting structures. The data presented here show that when this foundation is lacking and actin cytoskeletal organization is decreased, the nascent sprout is unable to achieve coordinated cell movement. ANXA2 is crucial for junctional maturation, which in turn is necessary for sprout maturation. As ANXA2 knock-down results in excess DPPC, the result is destabilized junctions, followed by hyper-initiation of sprouting. However, the lack of junctional reinforcement ultimately causes failed sprout maturation at later stages of the process, involving Arp2 recruitment.

## Data Availability

Full lipidomics data available from the corresponding author upon request.

## Acknowledgements

We would like to extend our thanks to Dr. Stanislav Vitha at the Texas A&M Microscopy and Imaging Center for helpful advice on imaging analysis. This project was funded by the Department of Molecular and Cellular Medicine, Texas A&M Health Science Center.

