## Supplemental Figures for "Annexin A2 modulates phospholipid membrane composition upstream of Arp2 to control angiogenic sprout initiation"

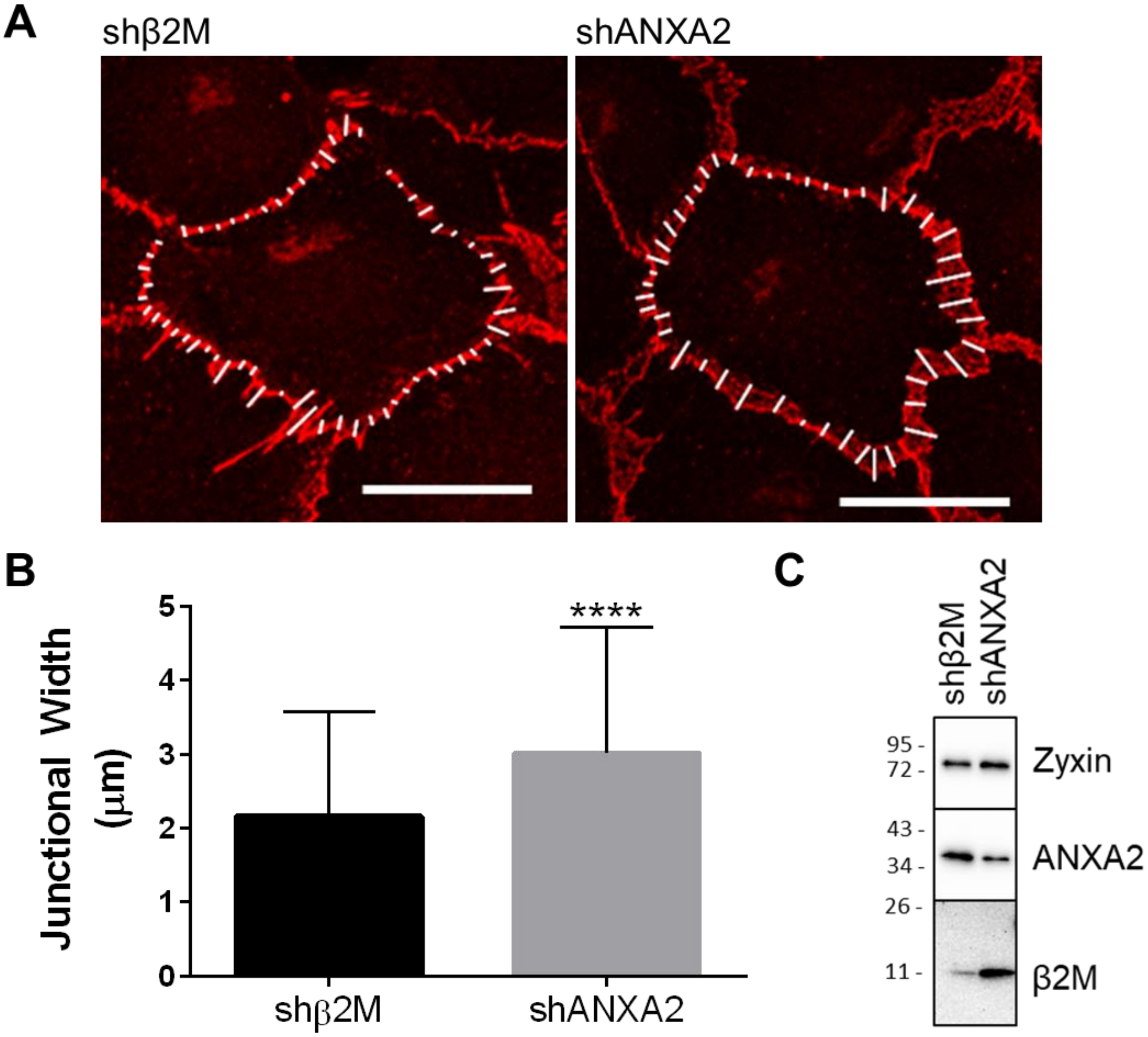


**Supplemental Figure 1** Junctional width increases with loss of ANXA2 during one hour of S1P activation. **A)** Immunofluorescence of VE-cadherin in shβ2M and shANXA2 cells, seeded onto collagen-coated coverslips and activated by S1P for one hour. White lines at junctions denote width measurement. Scale bar = 10 µm. **B)** Average junctional width (µm). N = at least 150 points per treatment. Student's t-test: ****, $p < 0.0001$. Experiment repeated three times with representative data shown. **C)** Western confirmation of knock-down. See figure 1G for quantification.

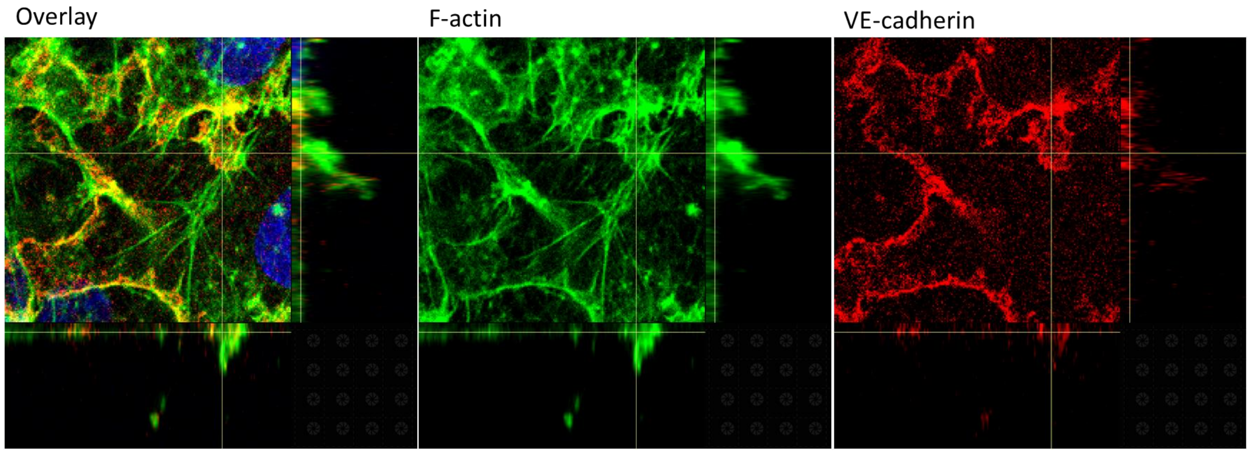


**Supplemental Figure 2** Orthogonal view of sprout initiation after 1 hour, generated by NIS elements. Yellow lines correspond with displayed X and Y slices.

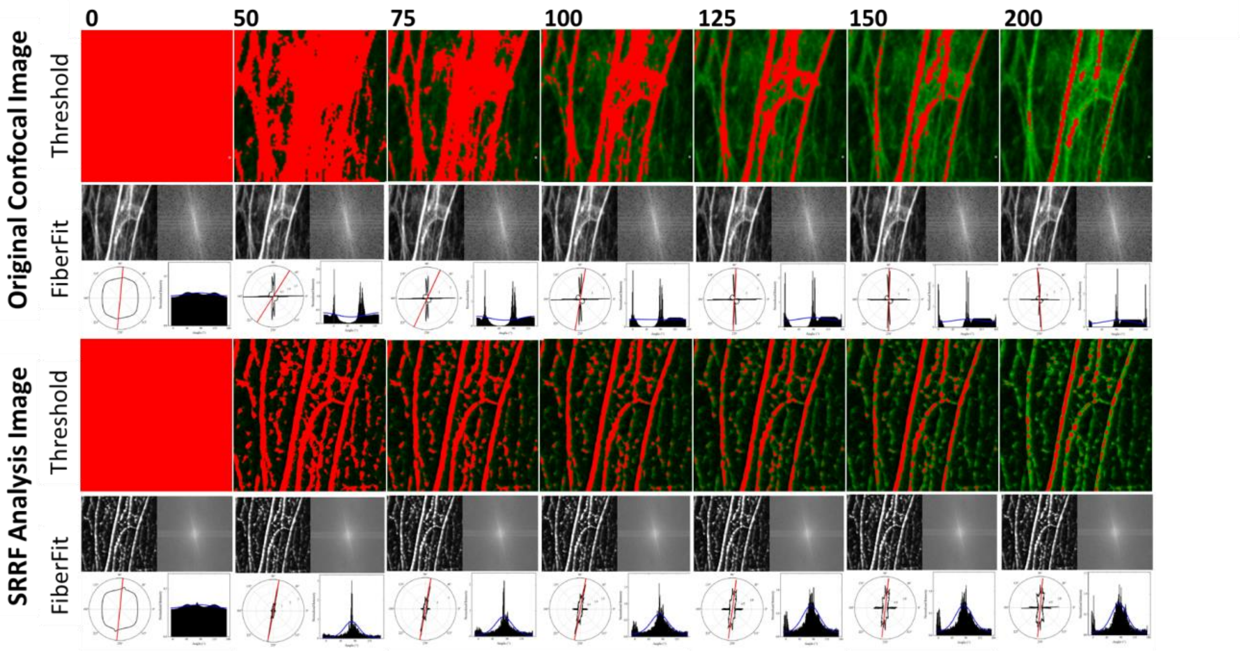


**Supplemental Figure 3** Benefits of SRRF analysis prior to FiberFit™. Cropped ROI of 100x confocal image taken at 1024 resolution. **Top)** Original confocal image at a given threshold (red denotes signal included for FiberFit Software™ analysis) and results output of that threshold. **Bottom)** Same field after SRRF analysis with different thresholds. Note: Threshold application with SRRF images provides a more representative output from FiberFit Software™ while preserving signal.

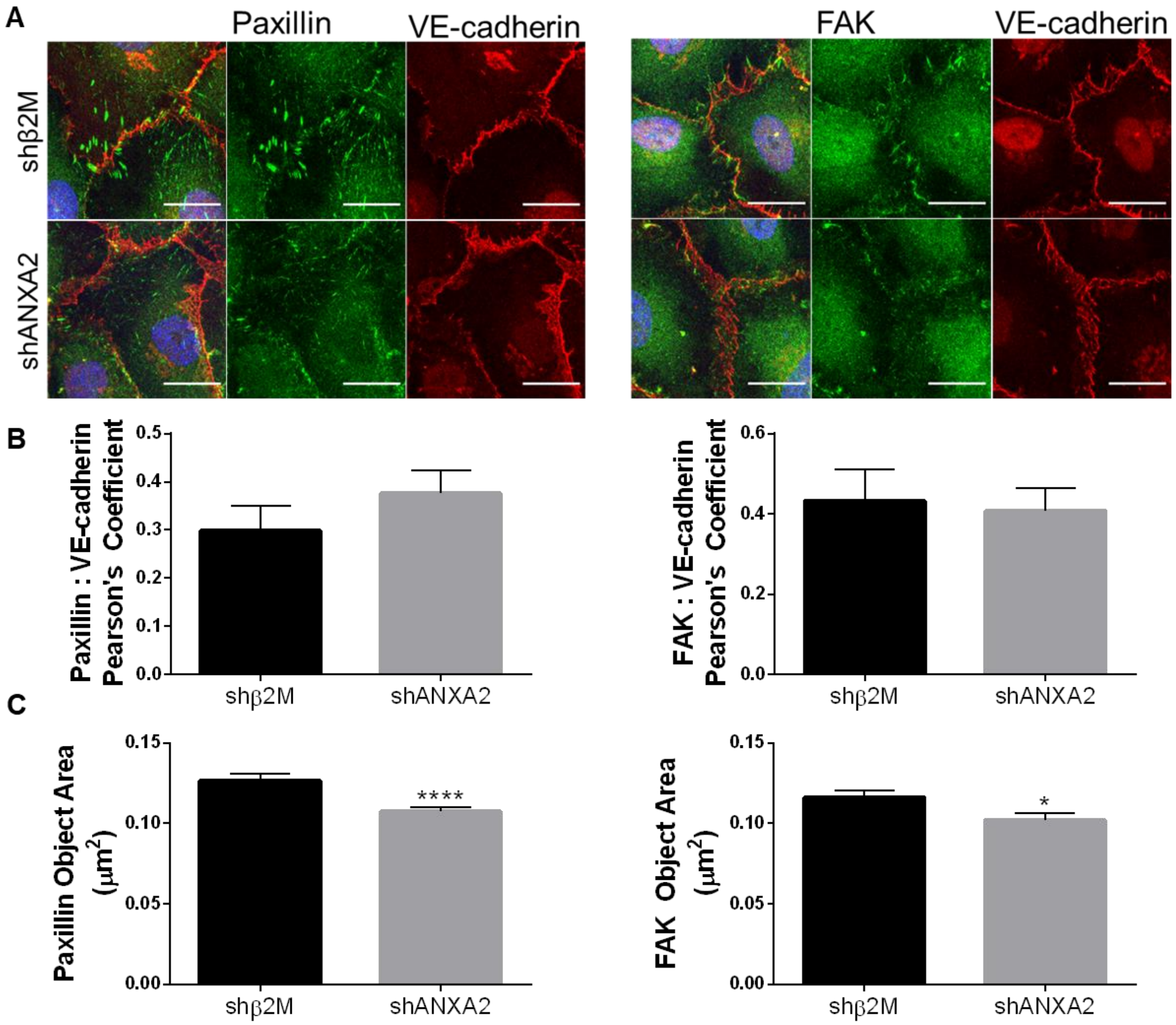


**Supplemental Figure 4** Focal adhesions show reduced sized with ANXA2 knock-down. **A)** Immunofluorescence of shβ2M and shANXA2 HUVEC after 1 hour S1P, then labelled for paxillin and VE-cadherin, or FAK and VE-cadherin. Scale bars: 10 μm. **B)** Quantified colocalization between FAK and VE-cadherin, or paxillin and VE-cadherin, as measured by Pearson's Coefficient. n = five fields per treatment. Student's t-test: p = Not significant. **C)** Quantified average object area using paxillin or FAK signal, denoting focal adhesions after applying a threshold. n > 2500 focal adhesions per treatment. Student's t-test: ****, $p < 0.0001$

for paxillin, *, $p < 0.05$ for FAK. All experiments repeated at least three times with representative data shown.

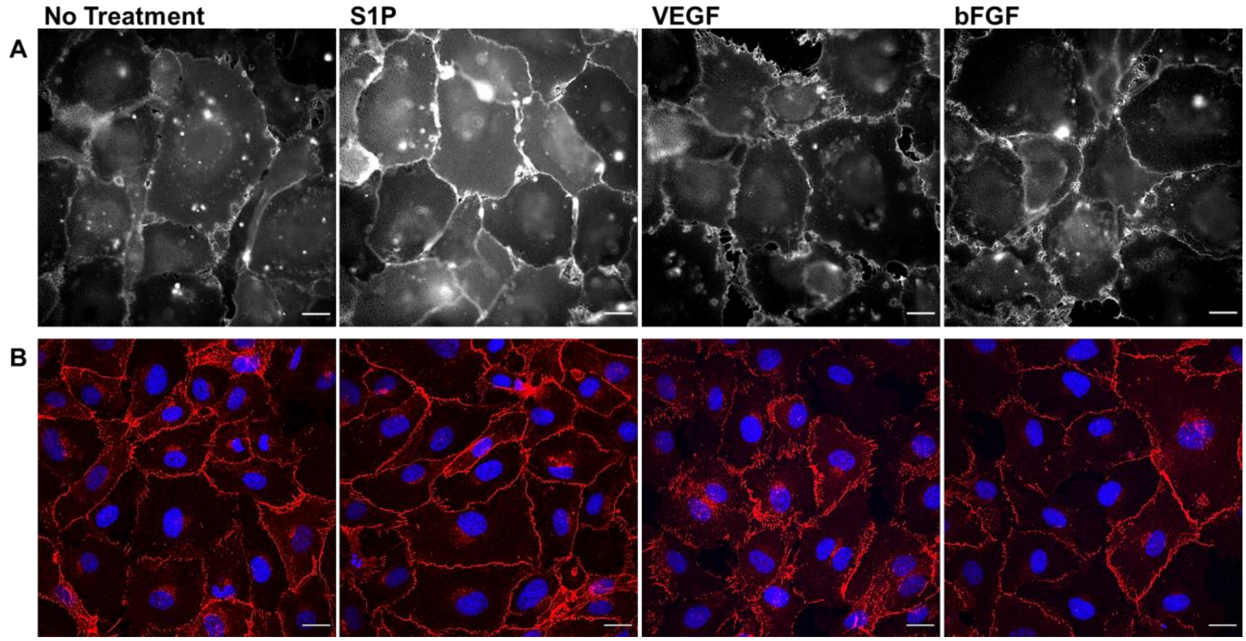


**Supplemental Figure 5** S1P enhances filipin III labeling of cholesterol at junctions but not VEGF or bFGF. HUVEC were seeded onto collagen-coated coverslips overnight, serum-starved, and treated for one hour with S1P, VEGF, or bFGF. **A)** Cholesterol localization as labeled by filipin III. **B)** VE-cadherin signal denoting adherens junctions. Note robust, linear signal at junctions in S1P-treated cells that is not present in cells with no treatment, VEGF, or bFGF. Scale bar: 10 μm.

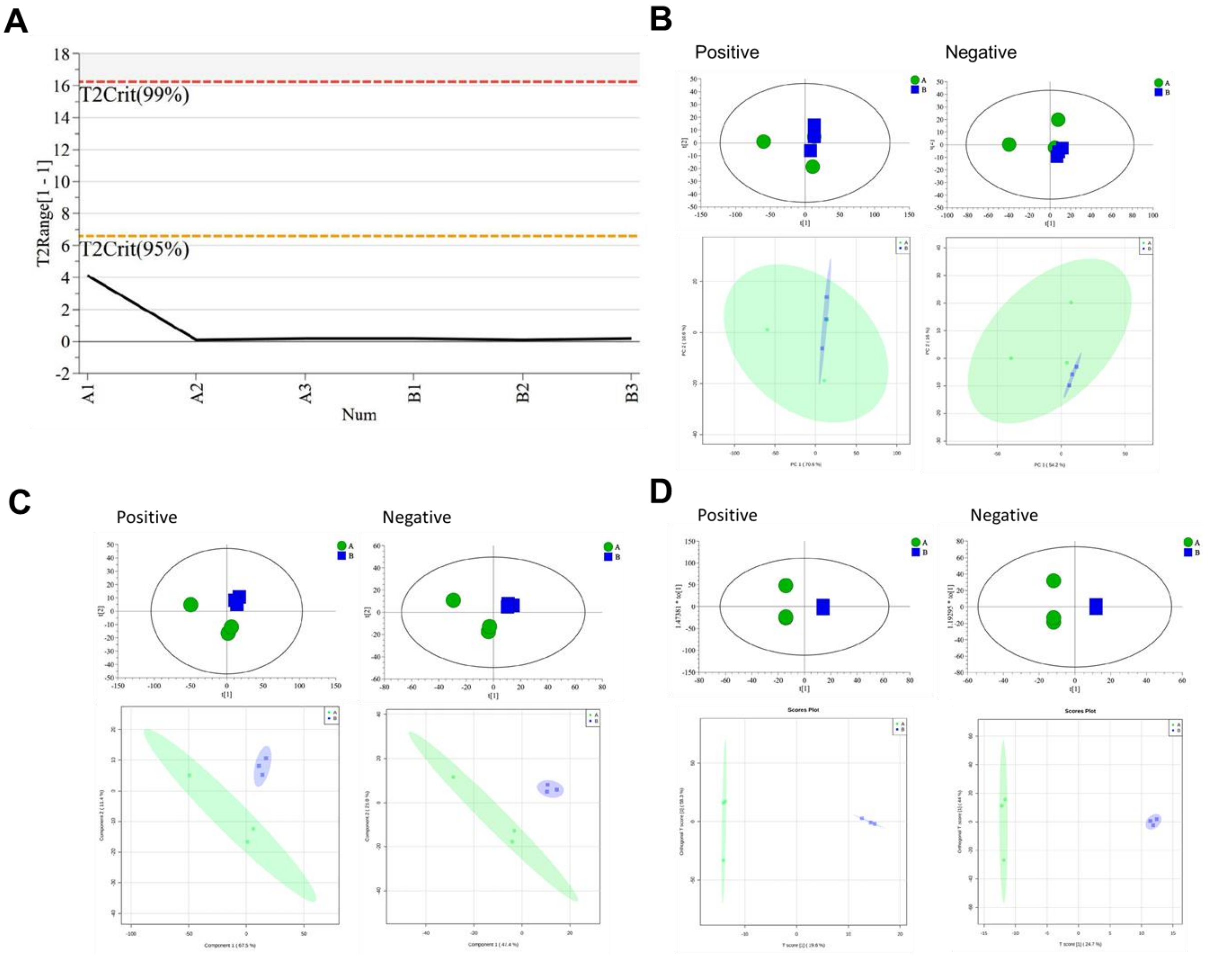


**Supplemental Figure 6** Quality control and modeling of LC-MS data via different clustering algorithms. **A)** Quality control of system stability prior to LC-MS analysis. A1-3 denote shANXA2 samples, B1-3 denote shβ2M samples. **B)** Clustering of samples by Principal Component Analysis. **C)** Clustering of samples by Partial Least Squares Discriminant Analysis. **D)** Clustering of samples by Orthogonal Partial Least Squares Discriminant Analysis.

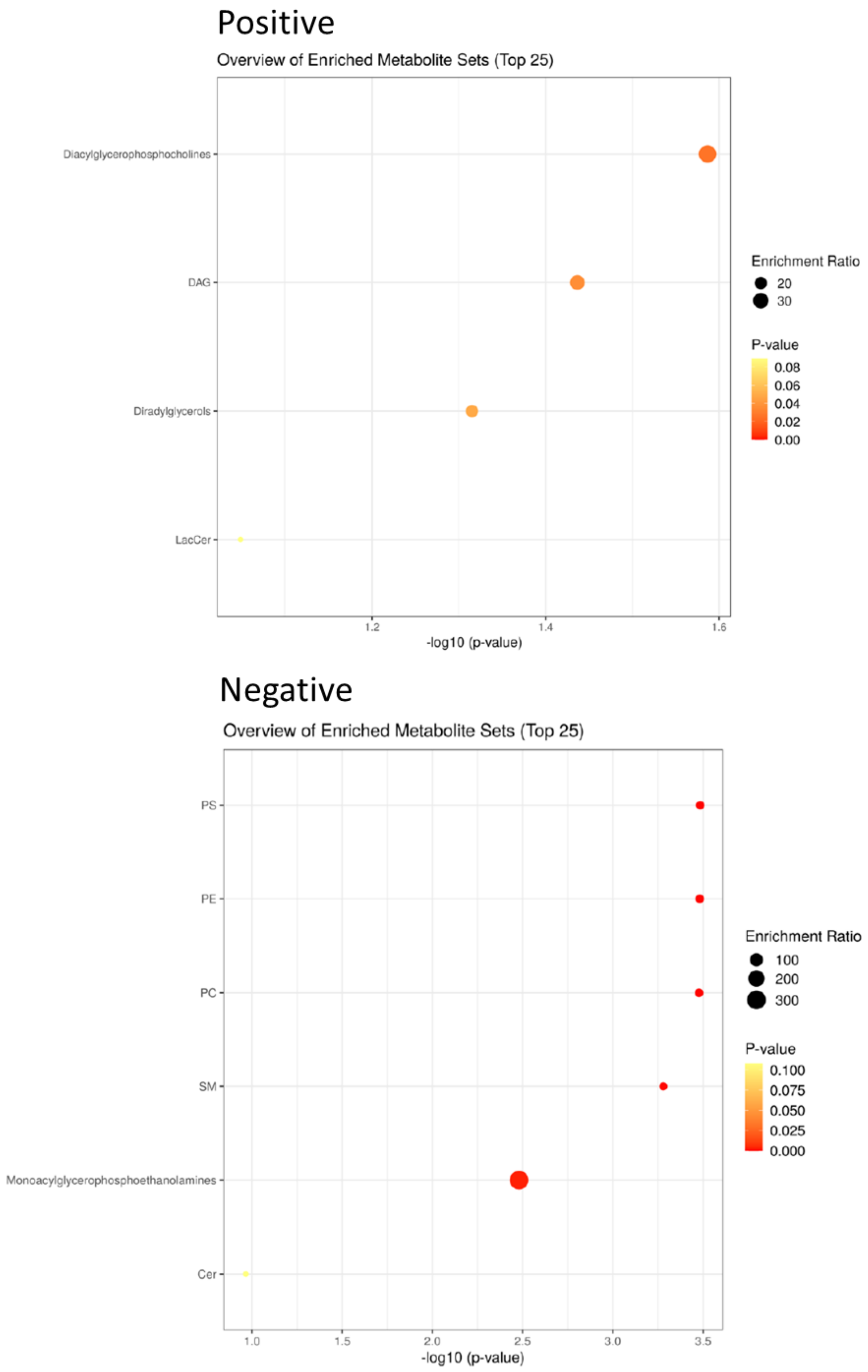


**Supplemental Figure 7** Pathways determined to be impacted by loss of ANXA2. Note: Plot is shβ2M relative to shANXA2.
